# A Swiftian Voyage from Brobdingnag to Lilliput: Freshwater Planctomycetes drifting towards the poles of the genome size spectrum

**DOI:** 10.1101/388082

**Authors:** Adrian-Ştefan Andrei, Michaela M. Salcher, Maliheh Mehrshad, Pavel Rychtecký, Petr Znachor, Rohit Ghai

**Affiliations:** Institute of Hydrobiology, Department of Aquatic Microbial Ecology, Biology Centre of the Academy of Sciences of the Czech Republic, České Budějovice, Czech Republic; Limnological Station, Institute of Plant and Microbial Biology, University of Zurich, strasse 187, CH-8802 Kilchberg, Switzerland

**Author notes:** Corresponding authors: Adrian-Ştefan Andrei &Rohit Ghai Institute of Hydrobiology, Department of Aquatic Microbial Ecology, Biology Centre of the Academy of Sciences of the Czech Republic, Na Sádkách 7, 370 05, České Budějovice, Czech Republic.

## Abstract

Freshwater environments teem with microbes. Currently, our apprehension of evolutionary ecology of freshwater bacteria is hampered by the difficulty to establish organism models for the most representative clades. To circumvent the bottlenecks inherent to the cultivation-based techniques, we applied ecogenomics approaches in order to unravel the evolutionary history and the processes that drive genome architecture in hallmark freshwater lineages from Planctomycetes phylum. The evolutionary history inferences showed that sediment/soil Planctomycetes transitioned to aquatic environments were, through processes mostly associated with reductive genome evolution, gave rise to new freshwater-specific clades. The most successful lineage was found to simultaneously have the most specialized lifestyle (increased regulatory genetic circuits; metabolism tuned for mineralization of proteinaceous sinking aggregates; psychrotrophic behavior) and to harbor the smallest genomes, highlighting a genomic architecture shaped by niche-directed evolution.

## Introduction

Planctomyces bacteria *(sensu* Woese et al.) [1] encompass one of the most enigmatic branches of the prokaryotic tree of life that have been brought into axenic culture [2]. This division, envisioned as a phylum [3], was thought to accommodate members that either rooted deeply in the bacterial line of descent [4] or paved the way to eukaryality [5, 6]. The obscurity surrounding the phylum arose decades ago when Nándor Gimesi described what he considered an unusual planktonic fungus (i.e. *Planctomyces bekefii)* in the eutrophic waters of Lake Langymanyos (Budapest, Hungary) [7]. However, this microbe which was later acknowledged to be of bacterial origin [8] was used to denominate the phylum Planctomycetes (Gr. adj. planktos wandering, floating; Gr. masc. n. mukês fungus; N.L. masc. n. Planctomyces floating fungus). The atypical morphology (e.g. microcolonial rosettes of cells joined together at the tips of their stalks) that misled Gimesi was found to be the norm for a phylum that accommodates bacteria with a vast array of shapes (from spherical and ellipsoidal to bulbiform), appendages (from spikes and bristles to stalks) [9–12] and outer membrane crateriform complexes [10]. Moreover, their puzzling appearance was found to be accompanied by a cell plan that seemed to diverge from the classical bacterial ‘Gram-negative’ one, due to the: i) apparent cytosolic compartmentalization [13], ii) lack of peptidoglycan (i.e. a hallmark of free-living bacteria) [14] and iii) presence of an endocytosis-like macromolecular uptake mechanism (a process universal among eukaryotes) [15]. The phylum’s peculiarities generally withstood genome-centric analyses, that in a way further deepened the knowledge gap by revealing the presence of a large ‘ORFan black hole’ (functional prediction for only 32–54% of ORFs) [16–18] and ‘giant genes’ [19] harbored by huge genomes (median genome size of sequenced Planctomycetes is 7.4 Mb in comparison to the more typical 3-4 Mb of other sequenced genomes). In light of recent research not only is the supposed ‘link’ to eukaryotes a product of convergent evolution [20], the endocytosis-like macromolecule uptake questionable [21] and cell plan an altered ‘Gram-negative’ one [22], the integration of genomic and structural data into an ecological framework also lags behind.

Consisting of two classes (i.e. Phycisphaerae and Planctomycetacia) that exhibit global ubiquity [23–27], the Planctomycetes phylum evaded extensive ecological characterization as a result of the inability to bring environmentally abundant representatives into axenic culture, or to access their genomic information [28]. In spite of their initial description in freshwater environments [2, 7], the majority of ecological and genomic studies were performed on marine ecosystems and seawater isolates [16, 18, 23, 29]. Although they represent one of the major prokaryotic groups in freshwater (with highly variable abundances from <1% up to 22%) [27, 30–32] and have been shown to have major roles in dissolved organic matter fractionation [33], our understanding of Planctomycetes is based on data derived largely from culture-based approaches [2, 34, 35], short reads analyses or/and hybridization-based techniques [27, 31, 32, 36]. While prone to primer coverage biases [37], the 16S rRNA gene-based studies pointed out that the abundant freshwater ribotypes do not have counterparts in culture and that their genomic diversity and ecological significance remains elusive [27, 32, 38]. Although in the light of recent research, Planctomycetes groups have been defined based on 16S rRNA gene relatedness (i.e. CL500-3, CL500-15 and CL500-37) and some are considered to be abundant in lakes and envisioned as hypolimnion specific [27], our apprehension of their ecology remains dim.

Here, we use ecosystem-scale taxonomic profiling (based on 298 metagenomic datasets), genome-resolved metagenomics (60 Planctomycetes genomes recovered from ten large metagenomic datasets) and spatio-temporal abundance patterns (using CARD-FISH) to elucidate the evolutionary history of lacustrine Planctomycetes, and to link their genome evolution patterns to their lifestyle strategies. In doing so, we not only characterized some of the most iconic freshwater bacterial lineages from an ecologic, genomic and metabolic perspective, but also broadened our view on genome evolution at large.

## Materials and Methods

The detailed description of the sampling sites, together with the experimental procedures and analyses performed are found in the Supplementary Materials and Methods.

## Results and discussion

### An aquatic Planctomycetes census based on short-read technology

To explore the taxonomic extent of aquatic Planctomycetes and to assess their contribution to prokaryotic community structure, we taxonomically profiled 298 metagenomic datasets derived from lacustrine (64 datasets), fluvial (36 datasets), freshwater sediments (40 datasets) and marine (158 datasets) habitats (Extended Data for a complete list). By making use of high spatial scale data (spread over four continents and along the Global Ocean) we show that Planctomycetes are ubiquitously present in aquatic habitats and sediments, where their contribution to prokaryotic assemblages varies (from absence to 13.13%) by environmental spatial heterogeneity (e.g. intralake; Figure 1) and to a lesser extent, habitat (with higher absolute abundances registered in freshwater habitats) (Figure 1, Supplementary Figure 1). For instance, the fluctuation in abundance (as assessed by the percentage of 16S rRNA gene reads), within prokaryotic community structure (e.g. Lake Zurich, samples collected on 13^th^ of May 2013; Figure 1), from scarcely present (0.11% rRNA gene reads in the epilimnion) to highly abundant (13.13% rRNA gene reads in the hypolimnion) pointed towards a niche, rather than habitat, preference. We observed that the taxonomic categories of aquatic Planctomycetes have a tendency to be more uniform within-than between-habitats (e.g. freshwater vs marine) (Figure 1), and that representatives of class Phycisphaerae form a major constituent of prokaryotic communities (up to 11.8% of total 16S rRNA reads) in lakes and reservoirs (e.g. Lake Zurich; Figure 1). Furthermore, we found that freshwater sediments harbored Planctomycetes communities with a higher richness (number of taxonomic categories) and evenness (relative abundance of taxonomic categories) than the aquatic habitats (Figure 1, Supplementary Figure 1), suggesting that they act as a ‘diversification hubs’ and may serve as starting points for adaptive radiations.

**Figure 1.**
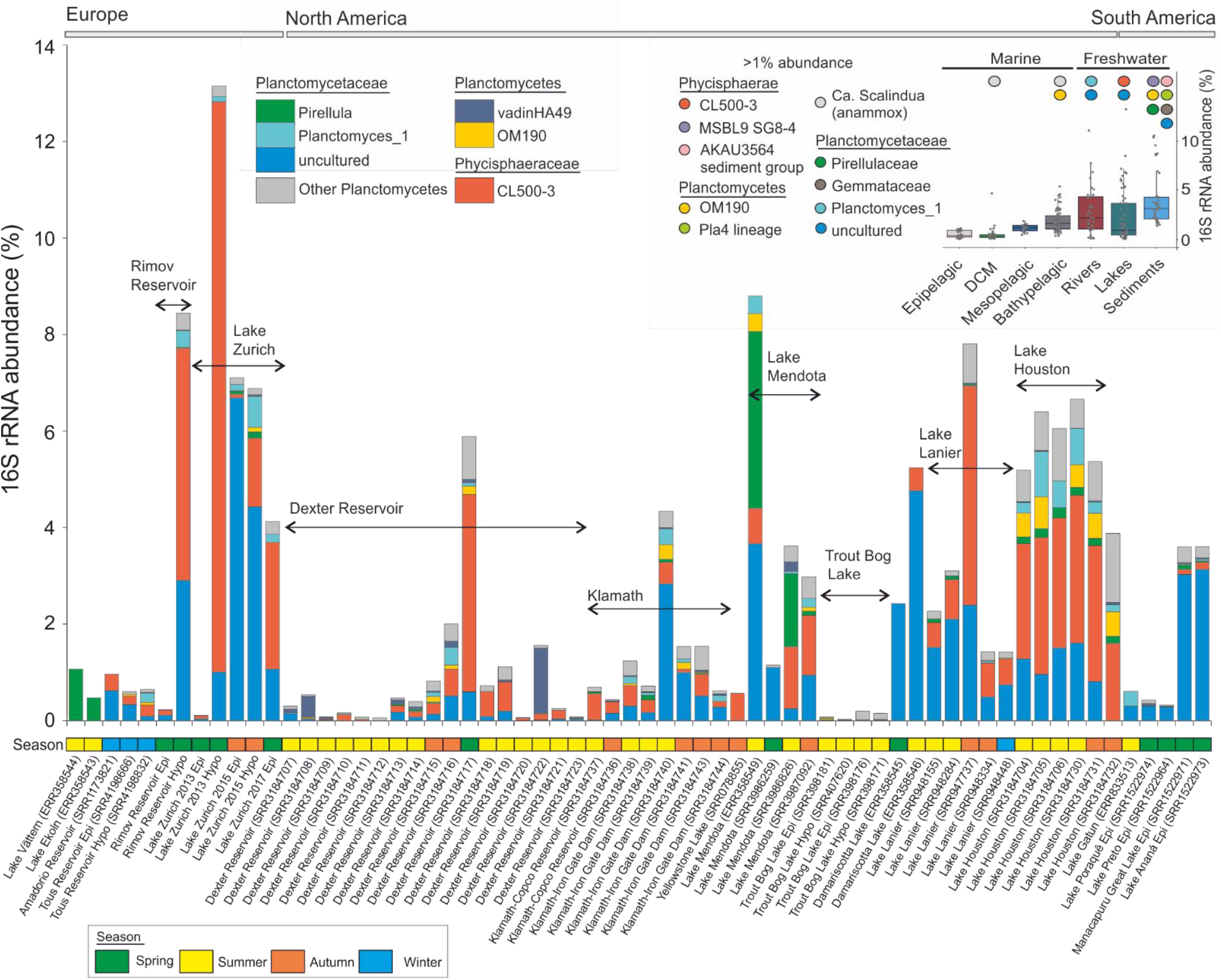
Taxonomic milieu of Planctomycetes phylum in worldwide lacustrine habitats. The figure depicts the SILVA SSU (Ref NR 99 128) classification of 16S rRNA gene fragments (as unassembled shotgun reads) retrieved from 64 freshwater datasets. The X-axis shows the taxonomic ranks and the geographic distribution of the sample collection sites, while the Y one indicates the percentage of Planctomycetes within the prokaryotic communities (as assessed by 16S rRNA genes abundance). The sample collection time, following a four-seasons breakdown, is indicated by colored boxes arranged along the X-axis. The SRA identifier for each metagenome is indicated in the parentheses that follow the habitat name. The figure’s inset (upper right panel) shows the contribution of Planctomycetes (as assessed by 16S rRNA gene abundance in 298 metagenomic datasets) to the prokaryotic communities present in aquatic habitats and freshwater sediments (64 lacustrine, 36 fluvial, 34 epipelagic, 46 deep chlorophyll maxima, 16 mesopelagic, 62 bathypelagic and 40 sediments). The colored circles highlight taxa that reached more than *1%* abundance within prokaryotic communities. DCM = deep chlorophyll maxima.

In spite of the broad intra-phylum diversity (as represented by 16S rRNA genes), the taxonomic breakdown revealed the presence of a freshwater Planctomycetes blueprint (Figure 1) largely characterized by the dominance of clade CL500-3 (Phycisphaerae) and a collection of ‘uncultured’ groups of the Planctomycetaceae (Planctomycetacia). Regardless of their wide environmental distribution, the dominant taxa were found not to relate to phylum’s cultured diversity, or even to appertain to groups under-represented in sequence databases (i.e. CL500-3 group consists of 40 sequences in SILVA’s SSU Ref NR 99 128 dataset, where it represents only ca. 1.6 *%* of the Phycisphaerae sequences). While ‘Planctomycetaceae uncultured’ represents an umbrella taxonomic category (composed of multiple polyphyletic clusters without any cultured representative), the CL500-3 forms a cohesive phylogenetic clade, described initially in the deep water column of the ultra-oligotrophic Crater Lake (hence the name of the group) [38].

### A fine-scale phylogenomic picture of freshwater Planctomycetes

The applied hybrid binning strategy (taxonomy dependent, using homology searches and taxonomy independent, using tetra-nucleotide frequencies and mean base coverages) allowed the recovery of high-confidence Planctomycetes-affiliated contigs, and their segregation into individual metagenome-assembled genomes (MAGs). The obtained MAGs were further assessed for completeness and redundancy based on the presence of ubiquitous single-copy genes (360 Planctomycetes-specific genes) and amino acid identity between multicopy ones (Supplementary Figure 2). After performing additional data curation (using anvi’o software) we obtained 60 MAGs (9 548 contigs; total length 123.76 Mb; average contig length 12.91 Kb) that simultaneously met our quality criteria (completeness ≥ 10%, contamination ≤ 10%, number of contigs ≤ 500), and had an average coverage depth higher than 5-fold over 90% of the nucleotides (ensuring for high-confidence base identification) (Extended Data, Supplementary Figure 2). To the best of our knowledge, the present dataset of 60 MAGs encompasses by far the highest amount of genomic information available for freshwater Planctomycetes (in contrast the 7 903 UBA genomes dataset contains only six freshwater Planctomycetes MAGs) [39].

We emphasize that the obtained 60 MAGs represent ‘genomic pools’ of Planctomycetes populations that share high sequence identity, and that they do not accurately reflect the genomic make up of specific clonal lineages. The alignment of short metagenomic reads to the MAGs showed that freshwater Planctomycetes typically consist of ecologically coherent and sequence-discrete populations (characterized by 98.5 – 100% sequence identity), that can experience both panmictic and clonal lifestyles. For instance, we observed that the population represented by the MAG TH-plancto1 was undergoing a selective event (at the time of sampling) which was on the way of producing a (nearly) clonal population (Supplementary Figure 3). On the other side of the spectrum, the ZH-13MAY13-plancto44 population was found to harbor highly panmictic gene pools (Supplementary Figure 3).

The evolutionary relations and the taxonomic ranks of the 60 Planctomycetes MAGs were investigated through gene- and genome-focused phylogenies. The topological backbone of the phylogenomic tree was supported by the phylogenetic one (i.e. using 16S rRNA – the most-adopted phylogenetic marker), and both methods reinforced a three-clade branching pattern comprising anammox planctomycetes and the two classes Planctomycetacia and Phycisphaerae (Figure 2, Supplementary Figure 4). All our 60 freshwater MAGs branched within these two existing classes, where they formed monophyletic groups that were usually divergent from the cultured and metagenomics-recovered representatives (Figure 2, Supplementary Figure 4).

We found that 22 of the MAGs share a common evolutionary lineage within class Phycisphaerae (red box, Figure 2), which (at the time of writing) comprises only three cultured non-freshwater species (i.e. *Phycisphaera mikurensis, Algisphaera agarilytica* – both isolated from a marine alga and *Tepidisphaera mucosa* isolated from a terrestrial hot spring) from which one genome is publicly available (that of *Phycisphaera mikurensis).* This phylogenetic cluster (comprising 22 MAGs) seems to form an ecologically coherent aquatic group together with the marine *P. mikurensis* that shares a common ancestry with the deeper branching sediment-dwelling representatives of the class (branches represented by the MAGs: Phycisphaerae bacterium SM23_33 and Uncultured Phycisphaerae; Figure 2). Hereinafter, we made use of 16S rRNA genes as 4 MAGs from the 22 were found to have 16S rRNA genes (Supplementary Figure 4) to anchor the phylogenomic Phycisphaerae clade (comprised of 22 MAGs, red box) into the larger gene-based bacterial taxonomy, and show that the MAGs fall within the CL500-3 clade (Supplementary Figure 4), the hallmark taxonomic group of lacustrine habitats (Figure 1). Thus, in this study, we managed to recover not just one of the largest number of Phycisphaerae MAGs (n=22), but also the most extensive genomic repertoire of an ecologically relevant and abundant freshwater bacterial lineage that so far completely resisted cultivation-dependent and –independent analyses. As indicated by the clade topology within the phylogenomic tree (e.g. statistically supported monophyletic lineage; Figure 2) and its congruence within 16S rRNA phylogeny (Supplementary Figure 4), we propose to designate a taxonomic category, to encompass this uncultured group, in accordance with the guidelines of Konstantinidis et al. [40]. Based on average amino-acid identities between the 22 MAGs (that registered values lower than 65 %; Supplementary Figure 5) [41] we suggest the creation of the family-rank Nemodlikiaceae (fam. nov.; Slavic, fem. n. p., named after Nemodliki, tutelary deities of water in Bohemian and Moravian mythology), to formally denominate the taxonomic group previously known from 16S rRNA data as the CL500-3 clade.

**Figure 2.**
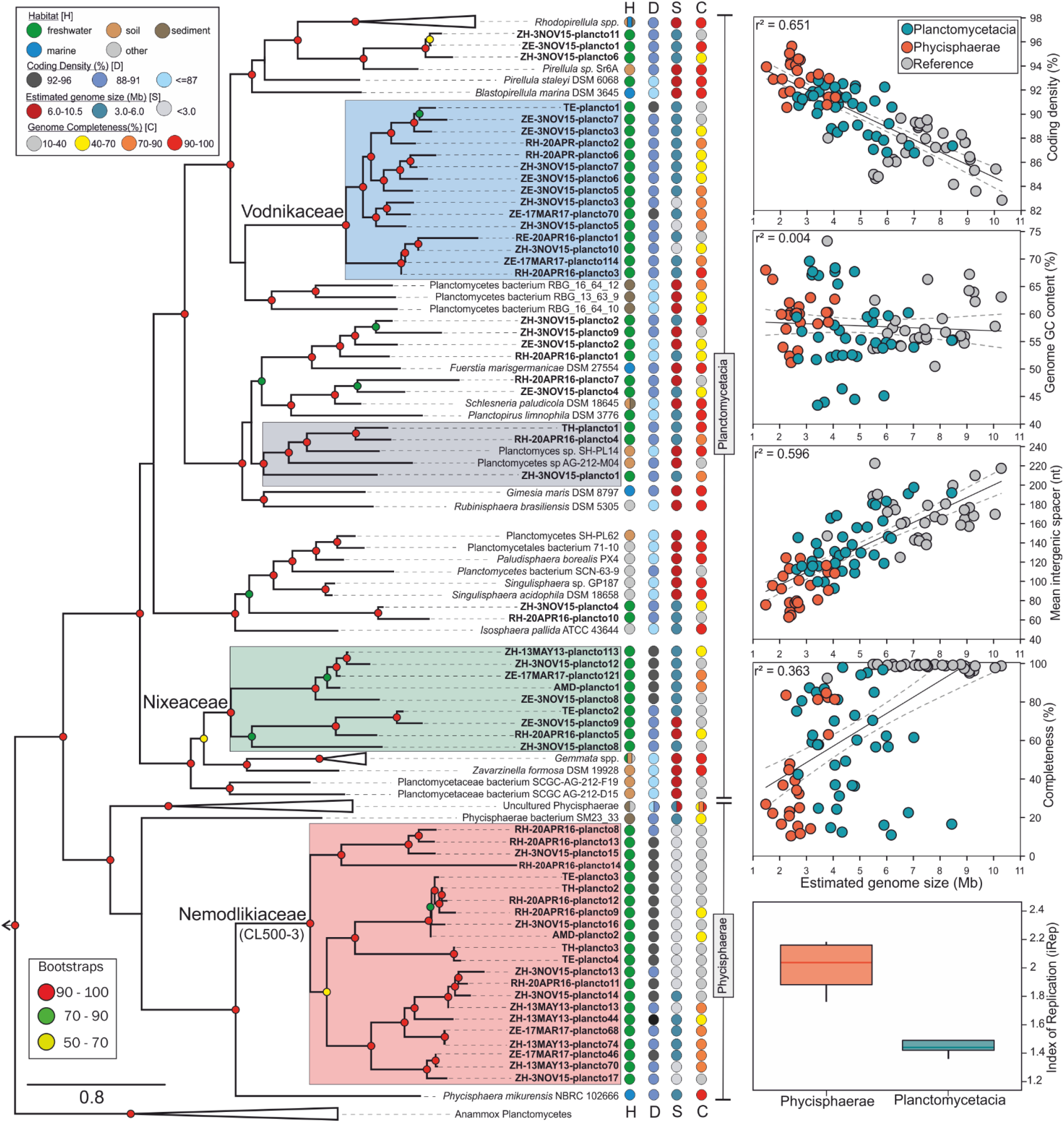
Phylogenomics of Planctomycetes phyla. The left panel shows accurate whole-genome phylogenies through a maximum likelihood (phylogenomic) tree inferred from 138 genomes (complete and partial). The topology of the tree emphasizes the major phylogenomic groups found in lacustrine habitats (for details regarding tree inference see Methods). The names of the 60 metagenome-assembled genomes (MAGs), obtained in this study, are highlighted in boldface, while the culture-derived genomes (references) and other available MAGs are depicted in italic and roman, respectively. The strength of support for internal nodes was assessed by performing bootstrap replicates, with the obtained values shown as colored circles (left legend}. Ecological data (i.e. habitat of origin=H) and genomic characteristics (coding density=D, genome size=S, and completeness=C) are indicated by colored circles for each branch in the tree (top left legend). The relations between the genomic characteristics (i.e. estimated genome size, coding density, GC content, mean intergenic spacer length, genome completeness) of MAGs (Phycisphaerae and Planctomycetacia MAGs; see vertical taxonomic delineators) and reference Planctomycetes (31 culture-derived genomes) are shown by linear regressions in the 4 insets present in the left part of the figure. The lowermost insert (right side) shows the iRep values for Phycisphaerae (n = 4) and Planctomycetacia (n = 9) MAGs.

The remaining 38 MAGs expanded the genomic representation of Planctomycetacia – the class that contains the bulk of cultured species (14 described genera at the time of writing) and considered (from a historical perspective) to encompass the planctomycetes *par excellence.* From them, 14 MAGs where found to form clusters affiliated to cultivated representatives (e.g. *Pirellula* spp., *Schlesneria paludicola, Planctomyces* spp., etc.), while the remaining 24 MAGs segregated in two coherent and divergent (from the other genomes and MAGs) groups within the class (Figure 2). The first one comprises 9 MAGs (green box, Figure 2) and branches in the proximity of *Gemmata/Zavarzinella* group, while the second (19 MAGs, blue box, Figure 2) shares an evolutionary ancestry with *Blastopirellula/Pirellula/Rhodopirellula* clade and appears to be phylogenetically more related to a sediment-derived MAG cluster (Figure 2). The 16S rRNA phylogeny showed that both of these clusters (green and blue boxes, Figure 2) fall under the umbrella rank “Planctomycetaceae uncultured” (i.e. Planctomycetaceae_uncultured G1 and Planctomycetaceae_uncultured G2; Supplementary Figure 4). As a consequence, based on within-group average amino-acid identity values (Supplementary Figure 5) we propose the creation of the families Nixeaceae (fam. Nov.; Germanic, fem. n., named after Nixe, aquatic being in Germanic folklore) (green box, Figure 2) and Vodnikaceae (fam. nov.; Slavic, masc. n., named after Vodník, mythical Slavic water spirit) (blue box, Figure 2) to accommodate the members of the 16S rRNA groups Planctomycetaceae_uncultured G1 and Planctomycetaceae_uncultured G2, respectively.

### Past and present of freshwater Planctomycetes explored by evolutionary genomics

The pattern of ancestry, divergence and descent (as showed by the phylogenomic trees, Figure 2 and Figure 3) indicated that Phycisphaerae and Planctomycetacia are sister lineages of a common ancestor which shared evolutionary relatedness with anammox planctomycetes (median genome size (MGS) 3.91 Mb, median intergenic spacer (MIS) 85 nt, median coding density (MCD) 86), bacteria that thrive at aerobic-anaerobic interfaces of sediment and water bodies [42].

**Figure 3.**
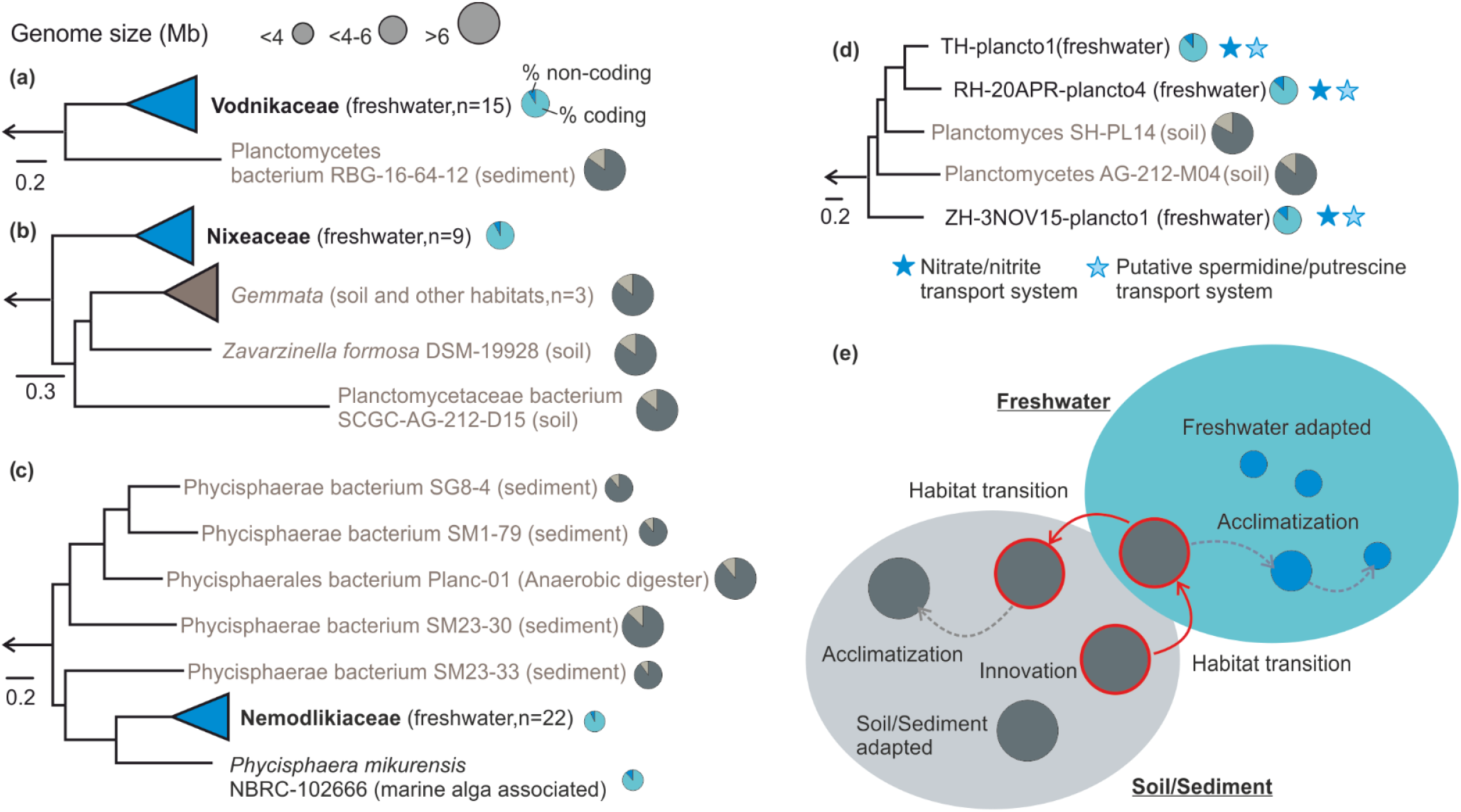
Phylogenomic subtrees (a, b, c, d) generated using maximum-likelihood methods and alignments of concatenated conserved proteins (54, 20, 206 and 315 proteins). The black coloured branches designate aquatic groups, while the grey ones their closest relatives (found in soil/sediments). The circular symbols, situated at the tips of the braches, are proportional with genome size and depict gene densities (within genomes). The blue stars highlight complete KEGG modules found in the freshwater genomes of the showed subtree (and absent in the soil-derived ones). The number of genomes present in the collapsed groups is specified in parenthesis, (e) Putative model of niche-directed genome evolution in freshwater Planctomycetes.

We observed that the deep evolutionary history of Phycisphaerae is intrinsically linked to a sediment-specific lifestyle, as the basal branch of the class was found to accommodate bacteria (MGS 6.13 Mb, MIS 76.5, MCD 89%) that live in estuarine sediments (‘Uncultured Phycisphaerae’ group comprising 4 MAGs; Figure 2, Figure 3). Both Nemodlikiaceae (MGS 2.6 Mb, MIS 43.5 nt, MCD 92.99%) and its sister lineage, typified by *P. mikurensis* (GS 3.81, MIS 97 nt, CD 88%) were found to be the descendants of an ancestor which underwent a habitat transition from sediments to an aquatic lifestyle (the node of the leaf Phycisphaerae bacterium SM23_33; GS 4.75 Mb, MIS 78, MCD 90%). Noteworthy, we observed that the adaptation to freshwater appears to be accompanied by a reduction in genome size (i.e. from 6.13 Mb in the deep branching sediment clade to 4.75 Mb in the sediment sister lineage of the aquatic branch and to 2.6 Mb in the freshwater Nemodlikiaceae) (Figure 3). The Vodnikaceae (MGS 4.17 Mb, MIS 73 nt, MCD 91.23%) and Nixeaceae families (MGS 4.71 Mb, MIS 59 nt, MCD 92.24%) were found to be related to lineages comprising soil/sediment planctomycetes that (compared to them) harbor considerably larger genomes (7.13 Mb MGS for Vodnikaceae sister lineage and 9.2 Mb for Nixeaceae sister linage) (Figure 3). Remarkably, these observations seem to be in agreement with the emergence of Planctomycetes at the dawn of the first terrestrial ecosystems [43].

Taken together, these observations point to the fact that freshwater Planctomycetes (as typified by the families Nemodlikiaceae, Vodnikaceae and Nixeaceae) may possess a sediment/soil ancestry, and that during adaptation to the freshwater environment underwent substantial genome downsizing. On the assumption that this hypothesis is accurate, we would expect that (within a ‘lower-rank’ taxonomic category) a transition from sediment/soil environments into freshwater will be accompanied by a decrease in genome size and *vice versa:* a freshwater-sediment/soil transition will be accompanied by an increase in genome size. Furthermore, the freshwater-specific foliage of such a phylogenetic cluster (depicting transitions from freshwater to sediment/soil and *vice versa)* would be characterized by bigger genomes (as compared to Nemodlikiaceae, Vodnikaceae and Nixeaceae), since they typify more recent habitat transitions. In line with the hypothesis, we identified in the phylogenomic tree a family-level clade (amino-acid identity within group 52.22 −66.86 %) (grey box Figure 2, Figure 3), in which the basal freshwater branch (ZH-3NOV15-plancto1, GS 5.3 Mb, MIS 82 nt, MCD 91.08%) is succeeded by a habitat transition to soil (Planctomyces sp. SCGC AG-212-M04: GS 6.93 Mb, MIS 107 nt, MCD 88%; Planctomyces sp. SH-PL14: GS 8.29 Mb, MIS 151 nt, MCD 83%) which is followed by a come-back to freshwater (RH-20APR-plancto4: GS 5.47 Mb, MIS 125 nt, MCD 87.06%; TH-plancto1: GS 5.27 Mb, MIS 107 nt, MCD 88.25%) (Figure 3). Furthermore, we observed that habitat transitions (from sediment/soil to aquatic environments and *vice versa)* are scattered throughout the evolutionary history of Planctomycetacia (Figure 2).

As the fitness of a prokaryotic cell (and its success in a heterogeneous environment) is generally considered to be dependent by its ability to modulate the gene expression patterns in response to environmental stimuli (temperature, pH, ionic strength, light, etc.), we investigated the distribution of signal transduction systems (STS) across the recovered Planctomycetes genomes. The 60 MAGs were grouped in a phylogenetic fashion (Nemodlikiaceae, Nixeaceae and Vodnikaceae) with the exception of 14 MAGs that did not generate discriminable freshwater specific clusters (i.e the 14 MAGs clustered together with cultivated representatives were grouped in ‘Planctomycetacia_diverse’). The inventory of sigma factors, signal transduction domains (histidine kinase A, Per-Arnt-Sim and GGDEF domains) and PP2C-phosphatases, revealed that Nemodlikiaceae harbored a surprisingly higher number of signal transduction pathways and genetic regulatory circuits per Mb of genome (in comparison to Nixeaceae, Vodnikaceae and Planctomycetacia_diverse) (Supplementary Figure 6). This inverse relation, in Nemodlikiaceae, (between STS/Mb and genome size) is unexpected since signal transduction systems and genome size are reported to positively correlate [44, 45]. Furthermore, in spite of harboring the largest genomes (MGS 5.37 Mb, MIS 109.5 nt, MCD 88.19%) within the 4 groups, Planctomycetacia_diverse was found to rank the lowest for GGDEF domains, and to respectively lack the histidine kinase A ones and PP2C phosphatases (Supplementary Figure 6). On the other hand, Planctomycetacia_diverse was found to contain the largest number of transposases (i.e. mobile genetic elements), which suggests an increased potential for genome plasticity and accelerated diversification through horizontal gene transfers and genomic rearrangements [46]. Taken together, the above observations suggest that signal transduction systems are critical components in the repertoire of freshwater Planctomycetes (that are retained in spite of genome shrinkage, increasing their genomic density) and may represent prerequisites for their survival and thriving in the lacustrine ecosystems. Moreover, we consider that the higher number of transposases found in Planctomycetacia_diverse may represent a genomic reminiscence that aided in habitat adaptation (Figure 3), and that their low numbers of signal transduction systems (together with their genome size and phylogenomic position) may be an indication of a more recent transition to freshwater environments in comparison to Nemodlikiaceae, Nixeaceae and Vodnikaceae.

### Freshwater Planctomycetes across space and time

Differential genome coverage was used to estimate the fraction of the Planctomycetes populations undergoing active DNA replication. By taking advantage of the coverage bias in actively replicating populations (as more sequences are recovered from the regions proximal to the origin, rather than the terminus of replication) and single time-point metagenomic sequences, we used the iRep algorithm [47] to infer *in situ* replication rates. We stress that a population in which the majority of the Planctomycetes are replicating the iRep value would be equal to 2. From the analyzed MAGs (13 MAGs that meet the iRep requirements: >=75% complete, <=175 fragments/Mbp sequence, and <=4% contamination) we inferred that on average 44% of Planctomycetacia and all the Phycisphaerae (Nemodlikiaceae) cells were undergoing replication at the time of sampling (Figure 2). Remarkably, the highest iRep values were registered for ZE-17MAR17-plancto46 (iRep = 2.18) and ZH-13MAY13-plancto70 (iRep = 2.15), MAGs belonging to the same species (Supplementary Figure 5) that were recovered at a four year-interval (from epilimnion and hypolimnion of Lake Zurich, respectively). The fact that the two MAGs had similarly high replication indexes, at different time points, suggests they represent a fast-growing genotype that is persistent and successful in the lacustrine habitats. Although, the low number of observations (4 for Phycisphaerae and 9 for Planctomycetacia) precludes generalization, it seems (from the available data) that the Phycisphaerae MAGs (i.e. Nemodlikiaceae) exploit more efficiently the available resources and thrive in the freshwater environments.

The biogeographic distribution of the 60 Planctomycetes MAGs was assessed in 64 lacustrine freshwater habitats scattered over three continents (Supplementary Figure 7). The results corroborated well with the 16S rRNA short-read taxonomic profiles and highlighted that, in general, the MAGs achieve higher ‘abundances’ in the habitat of origin, and scarcely few of them (e.g. AMD-plancto2, RH-20APR16-plancto14, ZH-3NOV15-plancto16, RE-20APR16-plancto1, TE-plancto2, ZH-3NOV15-plancto11) were well-represented in other European lakes. Considering that the majority of MAGs show a restricted geographic dispersal indicates that (in this case) the lakes’ low habitat connectivity supported a distributional pattern governed by a distance-decay relationship.

### A quantitative dimension of freshwater Planctomycetes revealed by CARD-FISH imaging

We made use of CARD-FISH technique to monitor the yearlong spatio-temporal distribution of Planctomycetes in Lake Zurich and Římov Reservoir throughout 2015. Hence, ten CARD-FISH probes were designed using the 16S rRNA gene sequences recovered from MAGs and additional public available sequences. Seven probes were constructed to target groups from which MAGs were available and another three were designed to quantify Planctomycetes groups that were found to be abundant in the metagenomic 16S rRNA gene pool but from which MAGs were not recovered (Supplementary Figure 4, Extended Data).

Nemodlikiaceae (class Phycisphaerae) numerically surpassed the other detected Planctomycetes with the exception of thermal stratification events when, in the warmer epilimnion, members of class Planctomycetacia prevailed (Supplementary Figures 8 and 9). We observed that the uniform abundance patterns of Nemodlikiaceae (Phycisphaerae) that were displayed within the water column during mixing in Lake Zurich, became skewed during stratification (Summer and Autumn distributions), when the group’s numbers declined in the epilimnion (Figure 4). A similar trend in spatial and temporal distribution was also detected in Římov Reservoir, where Nemodlikiaceae’s contribution to prokaryotic communities was at its lowest in the strata above the thermocline (Figure 4). Furthermore, we observed that Nemodlikiaceae i) maintained its high numbers in the surface strata long after the end of mixing events (6.6 – 7 % in April, Lake Zurich; 2.7 – 2.8 % in April, Římov Reservoir), ii) reached higher abundances in strata below the thermocline during stratification periods (Figure 4) and iii) registered seasonal peaks in abundances at low water temperatures (median temperature for seasonal peak in prokaryotic communities is 5.3 °C). By taking into account the above-mentioned effect of lake stratification-mixis cycles, we consider that Nemodlikiaceae are composed of habitat specialists (recording significant abundances in freshwaters, Figure 1) that show a trend towards psychrotrophic behavior *(sensu* Gounot) [48]. This is in line with previous studies that reported high abundances of this lineage in the deep and cold hypolimnion of several Japanese lakes [27, 31]. Nemodlikiaceae were found both free-living and attached to lake snow particles and had the smallest cell sizes of all analyzed Planctomycetes (ovoid shape, length 0.44 μm, width 0.31 μm; Figure 4, Supplementary Figure 10 and Extended Data).

**Figure 4.**
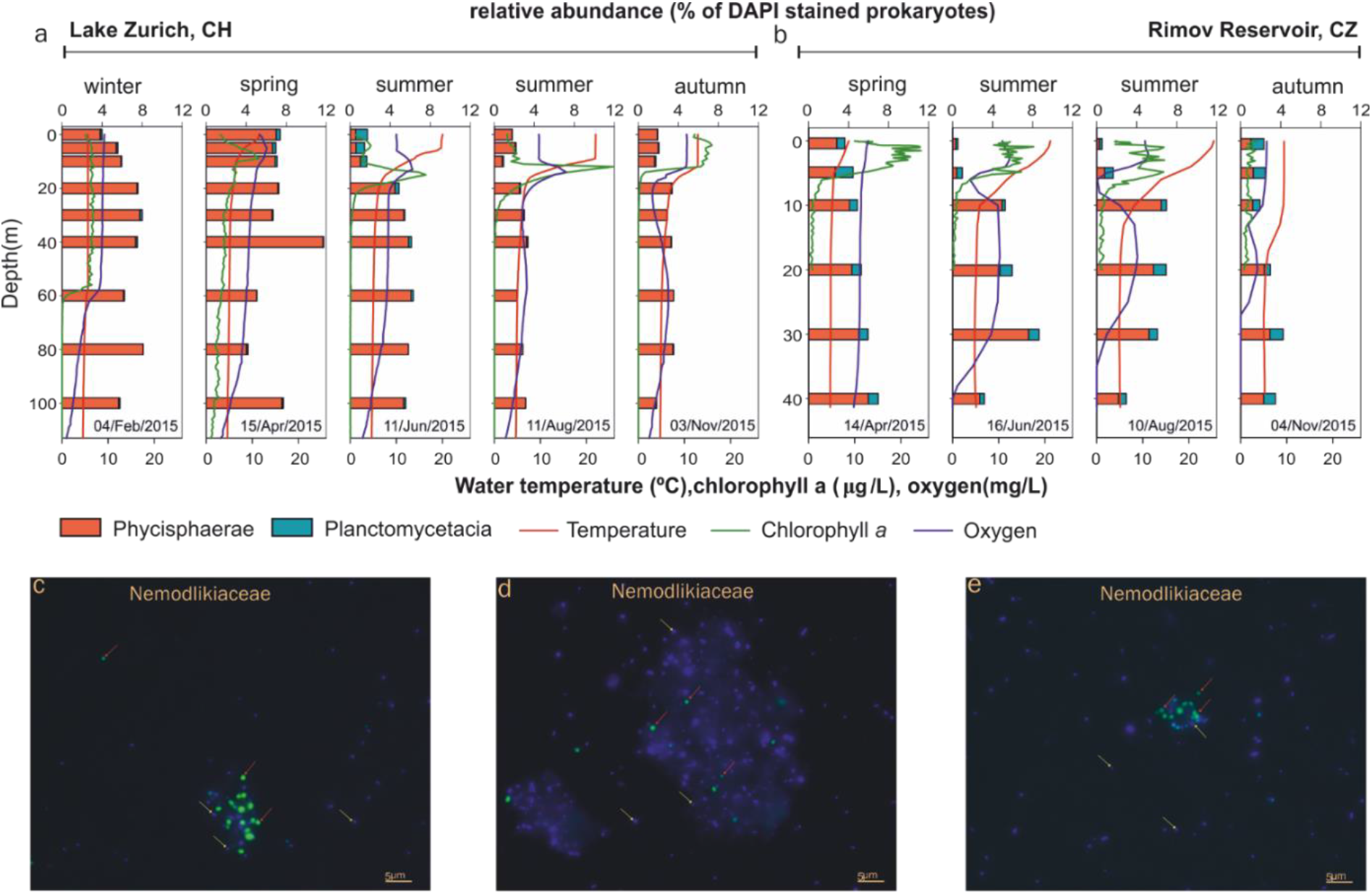
Spatiotemporal profiles of Planctomycetes relative abundance (horizontal bars), temperature (red line), chlorophyll a (green line) and oxygen (blue line) in Lake Zurich (a) and Rimov Reservoir (b) during 2015. The vertical axis shows the depth (m), within the water column, from which the samples were collected (9 for Lake Zurich and 6 for Rimov Reservoir). The upper x-axis shows the percent of Phycisphaerae (red bars) and Planctomycetacia (dark cyan) within the prokaryotic communities (estimated as the total sum of DAPI-positive cells), while the lower one displays the values for temperature, chlorophyll a and oxygen. The sampling date Is shown above the lower x-axis. The panels c, d and e display superimposed Images of CARD-FISH-stained Planctomycetes (class Phycisphaerae, family Nemodlikiaceae) and DAPI-stained prokaryotes. The red arrows point towards free-living and particle-associated Planctomycetes, while the yellow ones designate unhybridized prokaryotic cells. The scale bar Is 5 μm.

The overall contribution of Planctomycetacia to prokaryotic communities in Zurich Lake and Římov Reservoir was generally low (Figure 4). The most abundant group detected was Pirellula-like, which mostly maintained subunitary contributions in the prokaryotic assemblages (Supplementary Figures 8 and 9, probe pir-663). In both lakes, the contribution of Pirellula-like microbes peaked in the surface strata during stratification (1.4 %, 5 m, April, Římov Reservoir; 1.2 %, 0.1 m, June, Lake Zurich), concomitantly with phytoplankton blooms (mostly green algae) as inferred by high chlorophyll *a* values. These microbes were also found to colonize lake snow particles and were slightly larger than Nemodlikiaceae (rod-shaped, length 0.51 μm, width 0.39 μm, Supplementary Figure 10 and Extended Data).

All other CARD-FISH probes applied to samples of Římov Reservoir resulted in values close to or below the detection limit of the method (0.2% of DAPI stained cells, Supplementary Figure 9), therefore, they were not further analyzed for the Lake Zurich dataset. However, individual cells could be visualized and sized (Supplementary Figure 10), and their morphology generally indicated ovoid-to rod-shaped cells with lengths between 0.56 – 1.17 μm (Extended Data).

### Life in the lacustrine realm

Here, we explore the nature of Planctomycetes-environment interactions in a reductionist fashion centered on survival-reproduction strategies. Thus, our niche inferences stem from the means employed by bacteria to probe the physico-chemical landscape (e.g. respond to chemical gradients and uptake nutrients), since they typify ecological strategies (for increasing fitness) and allow general behavioral predictions.

We reason that Planctomycetes in lacustrine environments may adopt dual lifestyles (free-living and surface attached) since some lineages were microscopically observed to colonize particles (Figure 4) and they possess the capacity for both motility and adherence encoded in their genomic repertoire (Extended Data). Thus, while the presence of WspE-WspRF (all groups) and FlrB-FlrC (only Nemodlikiaceae) two-component systems may regulate surface affinities [49, 50], the flagellar apparatus (present in Nemodlikiaceae and 3 MAGs from Planctomycetacia_diverse) suggests directional swimming (Extended Data). The genome-scale metabolic reconstructions, performed on the 60 Planctomycetes MAGs, revealed a typical heterotrophic metabolism in which beta-Oxidation, the hexose monophosphate shunt and glycolysis (incomplete in Nemodlikiaceae) fuel the tricarboxylic acid cycle and oxidative phosphorylation (Extended Data). We observed that while the core metabolism was highly similar between Phycisphaerae and Planctomycetacia the substrate uptake capacity showed phylogenetic segregation. Thus, albeit glucose (through porin OprB), ribose, nucleosides and 3-phenylpropionic acid uptake was inferred to be common in both Planctomycetes classes, the preferences towards monosaccharides and organic acids showed group specificity. Accordingly, we found that the uptake of hexoses (i.e. L-rhamnose, L-fucose, D-glucose/D-mannose, D-gluconate), pentoses (i.e glycoside/pentoside/hexuronide and L-arabinose) and organic acids (glucarate, hexuronate, lactate, oxalate) was favored in Planctomycetacia, while D-fructose was preferred in Phycisphaerae (Nemodlikiaceae). Moreover, even though the uptake systems for amino acids (polar, basic and branched-chained groups) and oligopeptides were found to be common across both lacustrine classes, they were more abundant in Phycisphaerae (Nemodlikiaceae) (4.5 vs 2.07 transporter components/MB). Although the presence of ammonium/ammonia (the preferred nitrogen source for microbial growth) transport channels (i.e. AmtB) was a common feature within Planctomycetacia, they were not detected in Phycisphaerae, thus, Nemodlikiaceae may lack ammonium/ammonia uptake capacity. Additionally, the enzymatic repertoire necessary for pyrimidines and amino acids (i.e. methionine, leucine, tryptophan and histidine) degradation was present exclusively in Nemodlikiaceae, implying an important role of these compounds in fueling their metabolic machinery. We detected that the amino acid biosynthetic pathways were also distributed unequally among the phylogenetic groups and that while both classes were auxotrophic for methionine, phenylalanine and tyrosine, Nemodlikiaceae suffered additional impairments in the synthesis of threonine, valine/isoleucine, leucine and proline (Extended Data). Evidence for sulfate transport (through ABC transporters and Sul P permease family) was found to be present only in Planctomycetaceae, where the assimilatory reduction pathway was inferred to be complete. The widespread capacity to regulate (through PhoR-PhoB two-component system) the high-affinity acquisition of inorganic phosphate (through phosphate-selective porins OprO and OprP) pointed towards a phylogenetically conserved strategy among all lacustrine Planctomycetes (Extended Data).

*Inter alia*, we inferred that Planctomycetes cellular membranes are dotted by mechanosensitive channels (both large- and small-conductance) that could jettison cytoplasmic solutes during hypo-osmotic conditions, and that Nemodlikiaceae intriguingly decorate their external surfaces with sialic acids. Noteworthy, we detected the presence of five green-light rhodopsins (one in Phycisphaerae and 4 in Planctomycetacia, Supplementary Figure 11) and CO dehydrogenases (form II; Planctomycetacia) that may be involved in energy conservation through generation of proton motive force. We found that members of Planctomycetacia (Vodnikaceae, Nixeaceae and Planctomycetacia_diverse) have the capacity to enhance their fitness and increase their niche persistence by antagonizing their (non-self) neighbors with lethal, toxin injecting devices (i.e. type VI secretion systems) (Extended Data).

Surprisingly, we found that Nemodlikiaceae genomes encode cohesin/dockerin modules (signature-domains of cellulosome, Supplementary Figure 12) that did not fit in the established cellulosome model [51] since we found no evidence for their involvement in cellulose degradation. Thus, we hypothesize that Nemodlikiaceae may use instead a non-canonical cellulosome-like machinery to degrade polypeptides, for which we tentatively propose the term planctosome (Figure 5). This putative structure resembles in its complexity the mesophile’s simple cellulosome systems [52] and supports a new non-cellulosomal function [52, 53] for the high-affinity cohesin–xdockerin interactions. By combining homology, motif- and structure-based methods with protein domain co-occurrence (Supplementary Figure 13) we consider that the lamin tail domain-containing protein facilitates peptidases anchoring on the outer membrane and activation through the proprotein convertase P-domains, while the cohesin/dockerin containing one facilitates substrate binding through a hyaline repeat domain (Figure 5).

**Figure 5.**
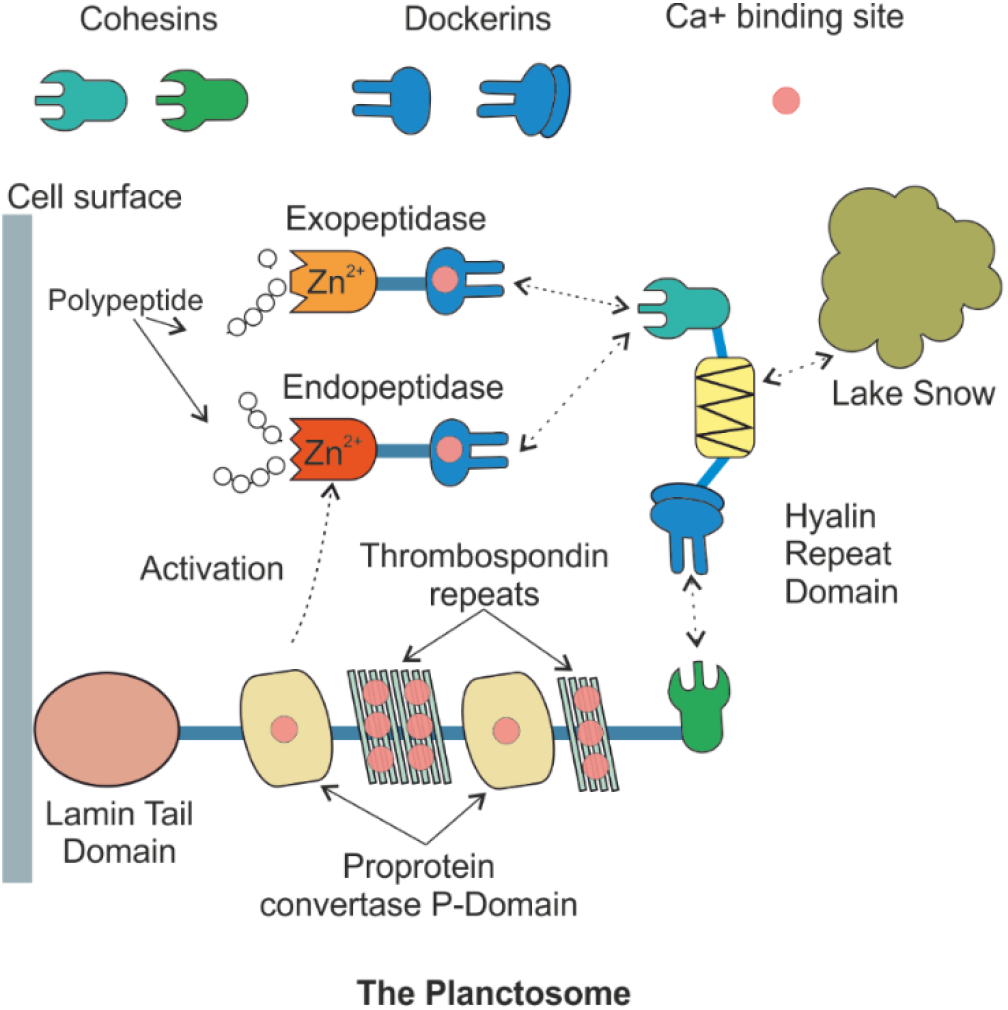
Hypothetical model of multiprotein complex (planctosome) involved in peptide degradation. The complex is tethered to extracellular membrane through a lamin A/C globular tail domain (LTD). The “anchoring” protein (2,210 aa) consists of a N-terminus signal peptide (26 aa) followed by the LTD, multiple proprotein convertase P-domains (PCD) divided by thrombospondin type 3 repeats (TSP), and a cohesin domain (CD). The “adaptor” protein contains a N-terminus signal peptide (26 aa), a dockerin domain (DD), a hyaline repeat one (HYRD) and a cohesin (CD). The “adapter” binds Zn^2t^-dependent endo-(M12B Reprolysin4-like) and exopeptidases (M14 carboxypeptìdase subfamily A) through Ca^2ł^-dependent cohesin-dockerin interactions.

## Conclusion

In line with the evolutionary history inference obtained by phylogenetic reconstruction, we suggest a scenario in which sediment/soil Planctomycetes transitioned to aquatic environments where they gave rise to new habitat-specific lineages (e.g. lacustrine-specific). Thus, we consider that freshwater Planctomycetes underwent a biphasic genome evolution process [54], in which an initial phase of innovation (characterized by acquisition of new genetic material) was succeeded by a phase marked by gradual loss of genetic material (in which metabolic versatility was sacrificed for efficiency). Thus, the innovation stage represented the trigger for habitat transition through acquisition of new gene sets (e.g. the freshwater genomes acquire gene sets that are absent in their sediment relatives, Figure 3), that eventually led to metabolic versatility and paved the way for niche expansions [55]. During this step of rapid gene acquisition, bidirectional habitat transition occurred (Figure 3), as the Planctomycetes had access to the genomic repertoire needed to survive in both soil/sediments and lacustrine environments. As the Planctomycetes entered the second (and much longer) evolutionary stage of acclimatization to freshwater, their genomes become smaller. We reason that the observed genomic shrinkage stems from the intrinsic trend towards gene loss, that governs bacterial evolution [54, 56–58].

While the performed large-scale taxonomic profiling (based on 298 metagenomic datasets; Figure 1 and Supplementary Figure 1) showed the existence of a lacustrine Planctomycetes blueprint, the *in situ* spatio-temporal abundance patterns and metabolic reconstructions pointed towards lineage-specific lifestyles. Thus, we observed that members of the Nemodlikiaceae (i.e. the hallmark lacustrine Planctomycetes lineage) exhibit psychrotrophic tendencies as they prefer to colonize the deeper and/or colder water strata, where they locate (by using signal transduction systems and flagella) and mineralize (through a highly-tuned metabolism) the nitrogen-rich sinking aggregates (lake snow, Figure 4). By contrast, Planctomycetacia (e.g. Vodnikaceae and Nixeaceae) showed preferences towards shallower and warmer water layers, where their versatile heterotrophic metabolism is fueled by phytoplankton-derived dissolved organic matter. Remarkably, the most successful lacustrine-specific Planctomycetes lineage (i.e. Nemodlikiaceae) had simultaneously the smallest genome sizes with highest coding densities and the most specialized lifestyle, suggesting niche-directed genome evolution. Thus, we consider that in Nemodlikiaceae the genetic drift fine-tuned their metabolic circuitry and decreased their genome size towards the minimum needed for efficient niche exploitation (selection of features necessary to colonize and utilize sinking aggregates; loss of biosynthetic pathways for molecules available in the niche).

Even though the composition of lake prokaryotic communities is usually depicted as being regulated by environmental filtering [59], the ways in which the niche shapes the bacterial genomic architecture and metabolic circuitry was previously unexplored. Here, we investigate the niche-genome interactions and show that: i) freshwater Planctomycetes bear in their genomes not only the marks of their ancestry but also the signatures of their lifestyle strategies, ii) substrate generalists (Nixeaceae and Vodnikaceae) maintain larger genomes than their more specialists counterparts (Nemodlikiaceae) and that iii) niche indirectly imposes constraints on the genome size through modulating the number of genes that could be lost (through genetic drift). By corroborating our results with recent phylogenetic reconstructions of abundant freshwater bacterial lineages (i.e. Betaproteobacteria and Verrucomicrobia) [60, 61] we consider that the above-mentioned evolutionary path in which ancient soil/sediment transitions are steered by the niche towards genome reduction may be wide-spread in freshwater ecosystems.

## Acknowledgements

The authors thank E. Loher and T. Posch for help with sampling of Lake Zurich and S. Neuenschwander for help with metagenomic library preparation for Lake Zurich samples. A-Ş.A was supported by the research grants: 17-04828S (Grant Agency of the Czech Republic) and MSM200961801 (Academy of Sciences of the Czech Republic). RG was supported by the research grant 17-04828S (Grant Agency of the Czech Republic). MM was supported by the Postdoctoral program PPPLZ (application number L200961651) provided by the Academy of Sciences of the Czech Republic.

## Supplementary Figures

**Supplementary Figure 1.**
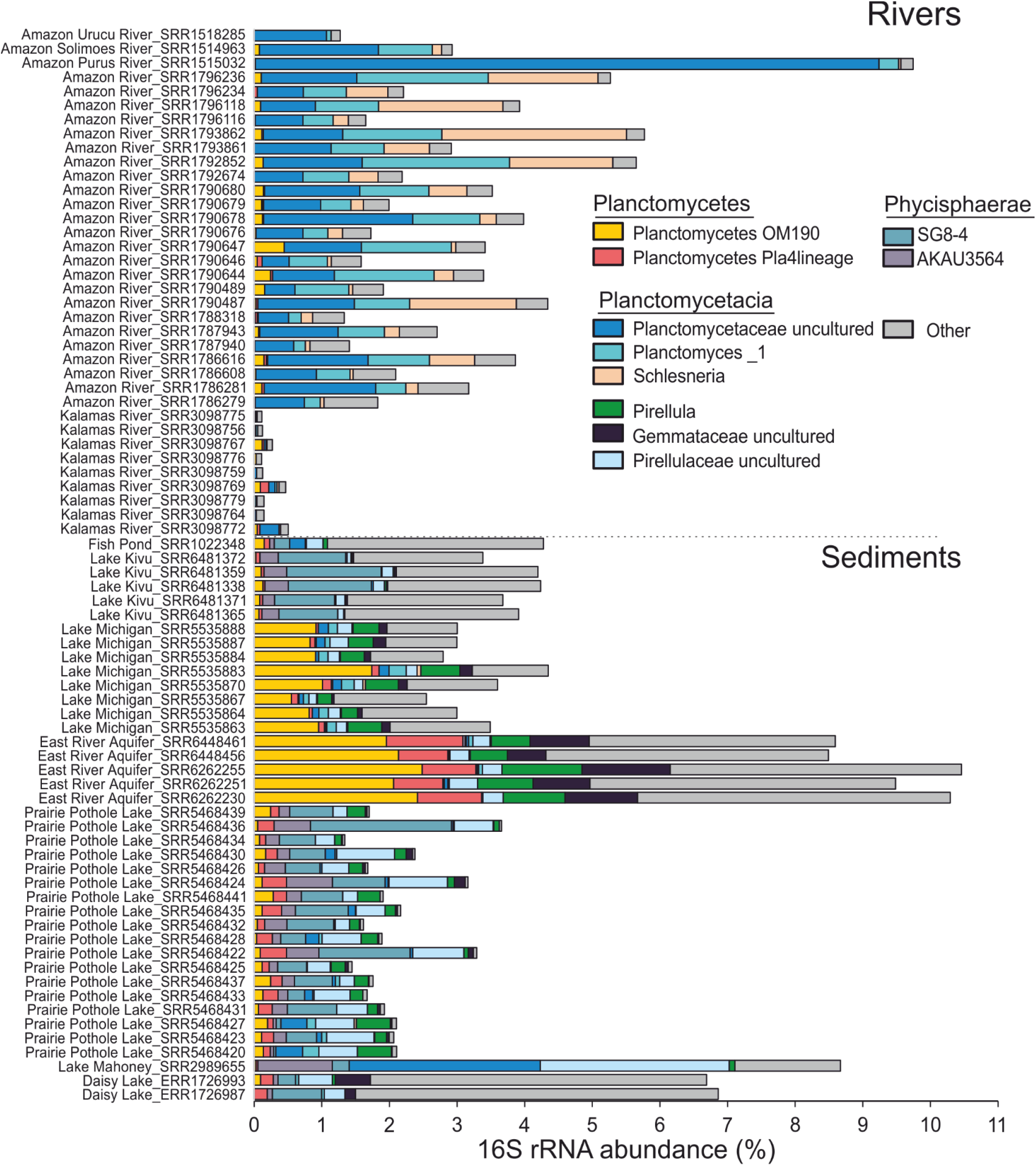
Taxonomic profile of Planctomycetes found in riverine and sediments ecosystems. The figure depicts the SILVA SSU (Ref NR 99 128) classification of 16S rRNA gene fragments (as unassembled shotgun reads) retrieved from 76 metagenomic datasets (36 from rivers and 40 from sediment samples). The X-axis indicates the percentage of Planctomycetes within the prokaryotic communities (as assessed by 16S rRNA genes abundance), while the Y-one shows the collection sites and their respective SRA identifiers.

**Supplementary Figure 2.**
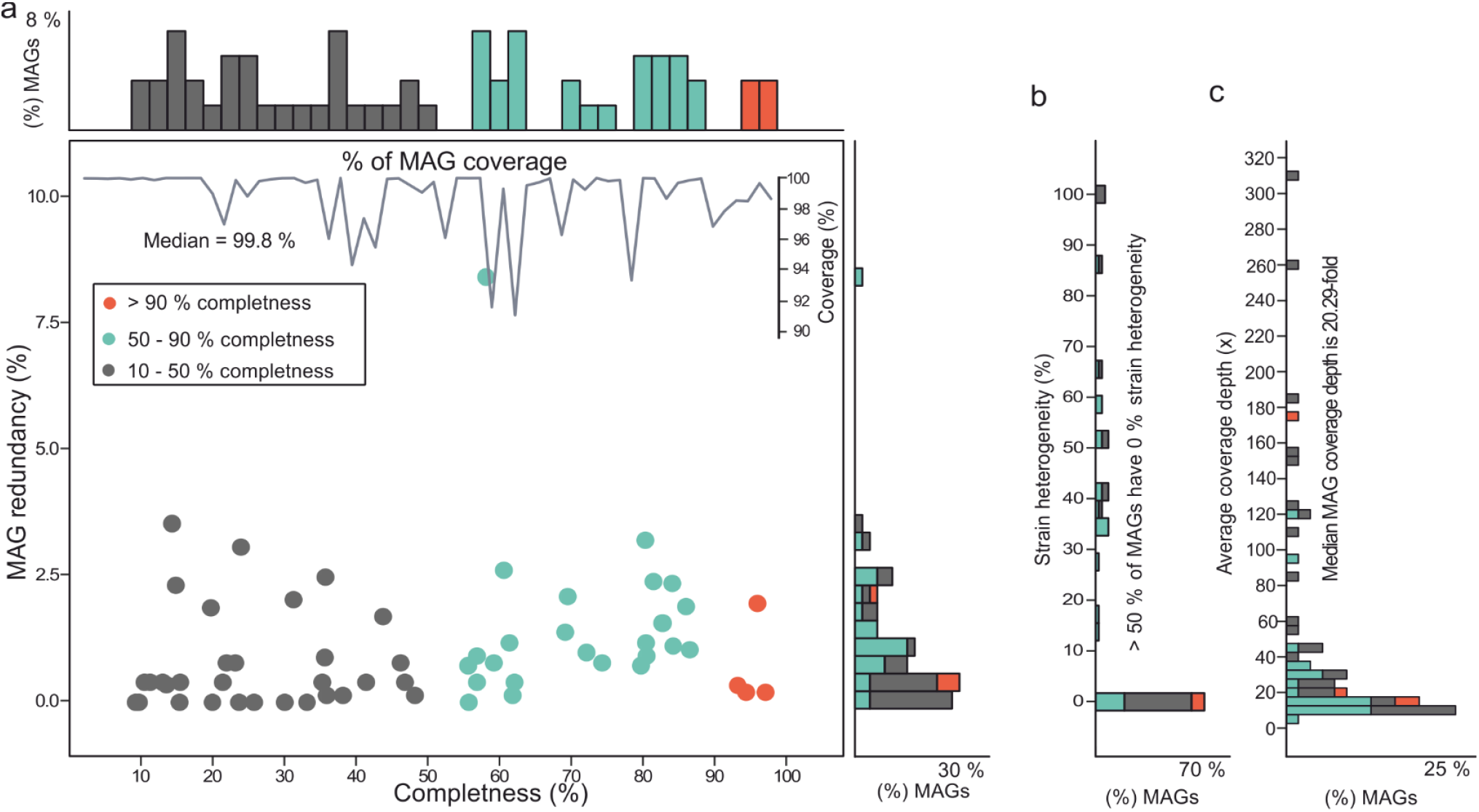
Metagenome-assembled genomes (MAGs) quality assessments. The figure shows the levels of completeness, redundancy and genomic coverage for each 60 Planctomycetes MAGs (panel a). The histograms situated along the X and Y axes depict the percentages of genomes that have varying levels of completeness and heterogeneity. The panels b and c, show MAGs heterogeneity and average coverage depth (from the metagenome of origin).

**Supplementary Figure 3.**
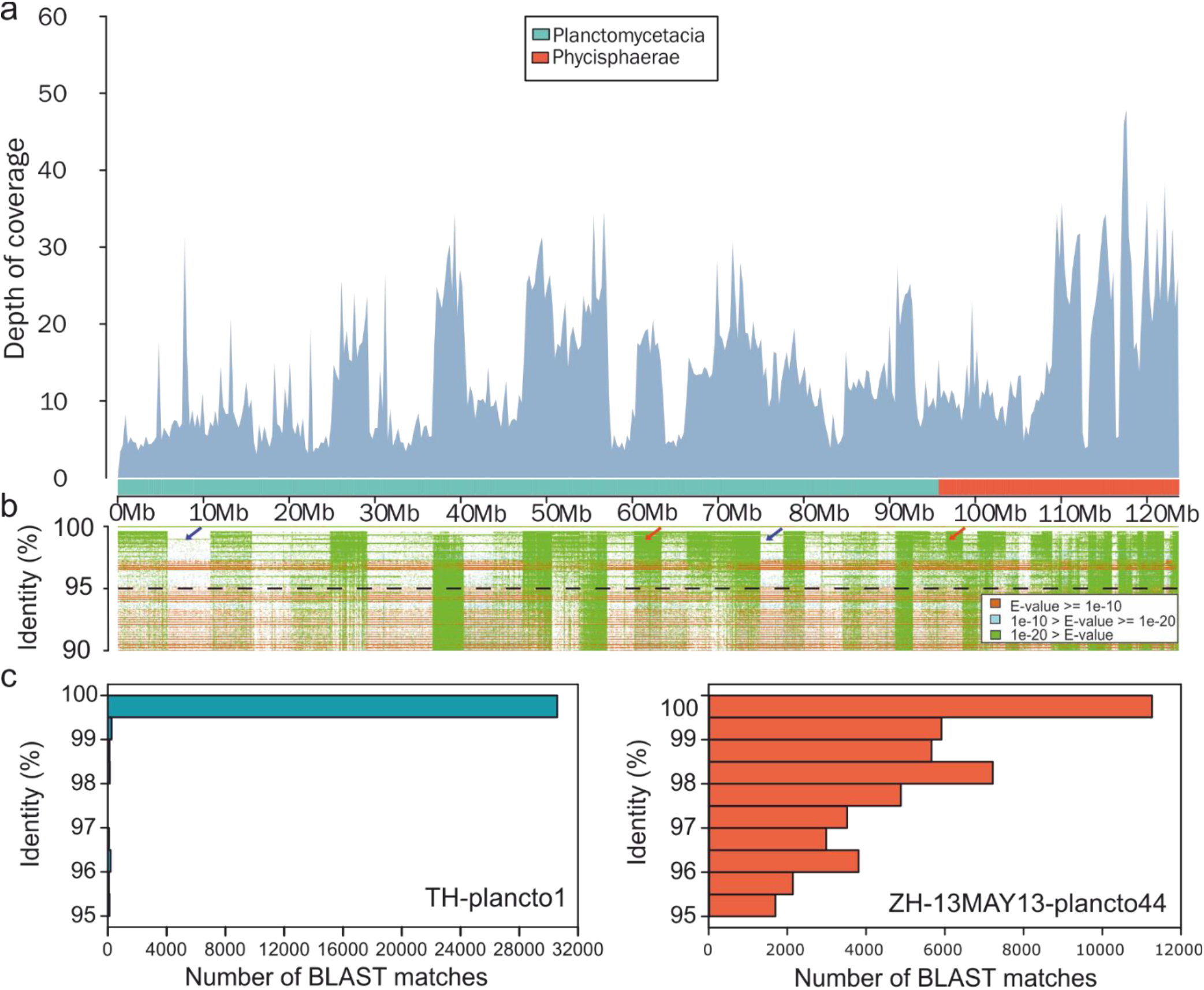
Planctomycetes MAGs recruitment plots. The panel a) shows the recruitment pattern of 180 million metagenomic reads (subsampled from the environments from where the MAGs where recovered) against the concatenated genomes of 60 MAGs. The ×-scale axis shows the total genomic length, while and the Y-one shows the coverage depth. The coloured bar situated above the X-axis indicates the taxonomic affiliation of the respective DNA fragment (dark cyan for Planctomycetacia, and red for Phycisphaerae). Panel b) shows the alignment identity (%) between metagenomic reads and the MAGs’ nucleotide sequences. Individual reads are coloured based on E-value scores (see legend in the left part of the panel). Red arrows indicate regions with high intra-population diversity, while blue arrows point towards ‘sequence-discrete’ populations. The histograms in panel c) show the number of reads (from 20 million reads subsets) (X-axis) that have identities > 95% with Planctomycetes MAGs. The MAG TH-planctol is a representative of ‘sequence-discrete’ species, while ZH-13MAYl3-plancto44 of high diversity one.

**Supplementary Figure 4.**
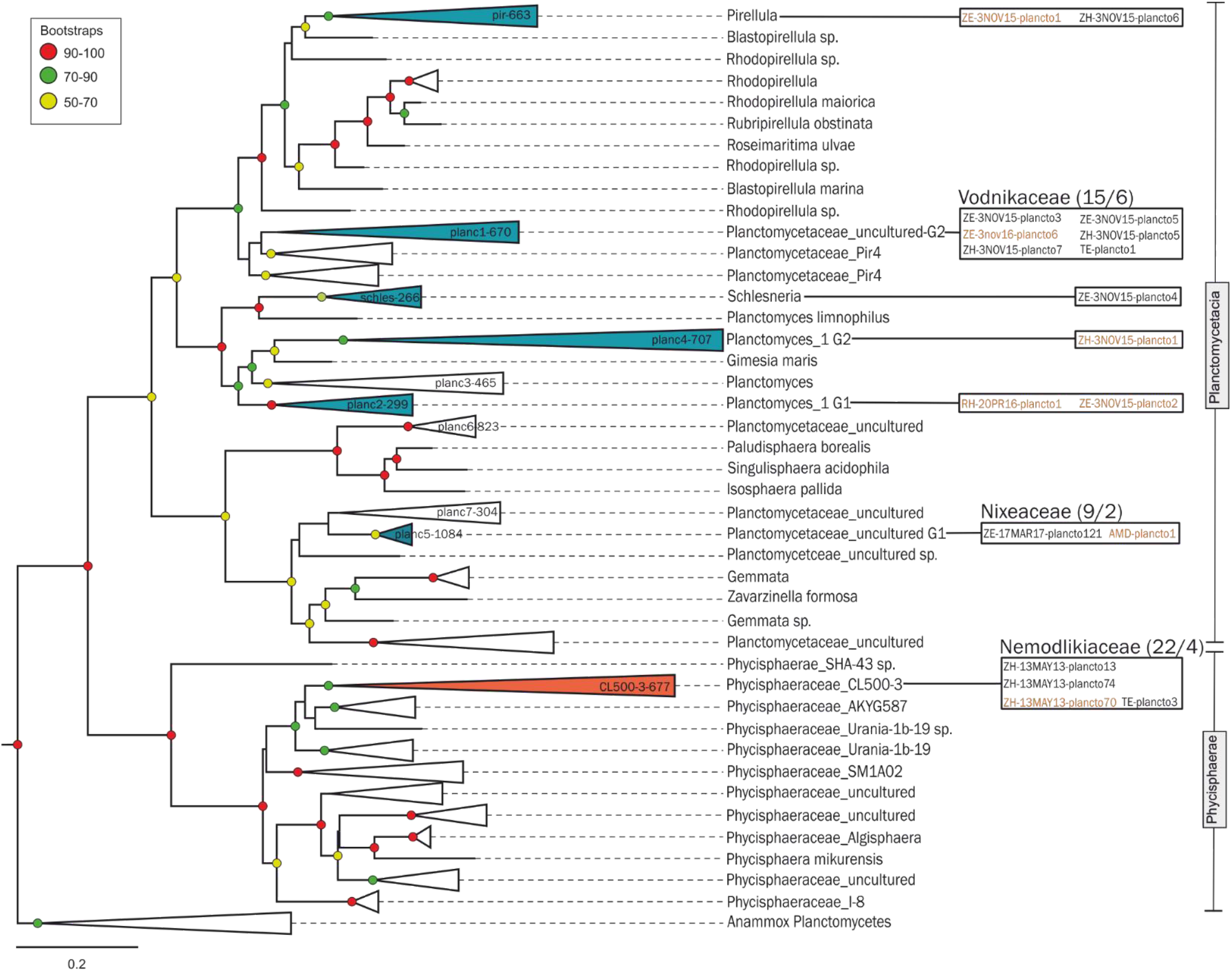
Maximum likelihood 16S rRNA phylogenetic trees. The names of the collapsed branches indicate the CARD-FISH probe that targeted the respective group. The left panels indicate the MAGs (from the respective clades) that had 16S rRNA gene sequences. The 16S rRNA gene sequences recovered from the MAGs with red colored names were used for CARD-FISH probe design. The strength of support for internal nodes was assessed by performing 100 bootstrap replicates, with the obtained values shown as colored circles (top left legend).

**Supplementary Figure 5.**
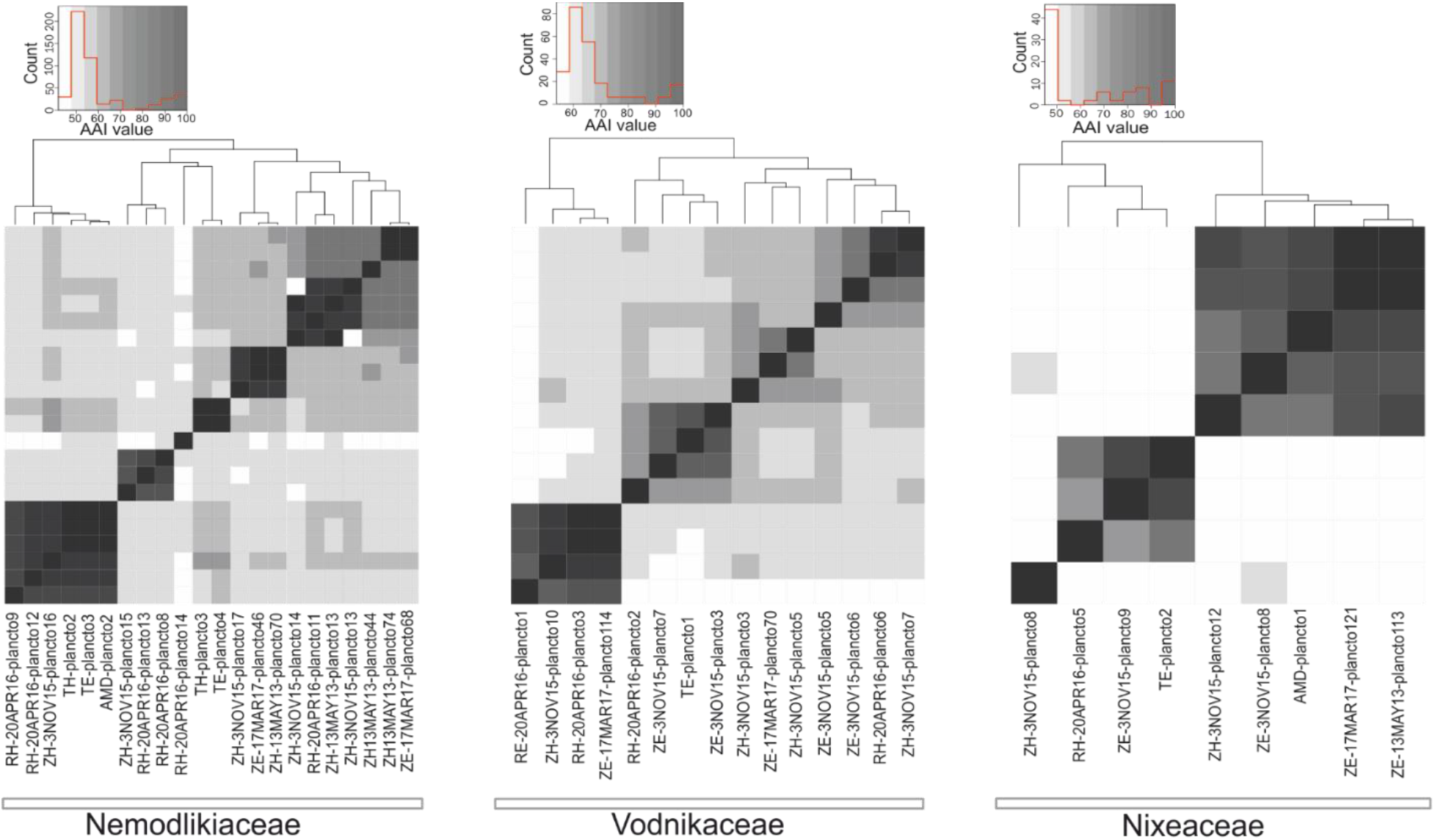
Heat maps of average amino acid identities between MAGs belonging to Nemodlikiaceae, Nixeaceae and Vodnikaceae families. The dendrograms positioned above the graphs show hierarchical clustering relationships between related MAGs. The names of the MAGs, and to the belonging phylogenomic groups, are shown undereach heat map. The upper panels display the color key histograms.

**Supplementary Figure 6.**
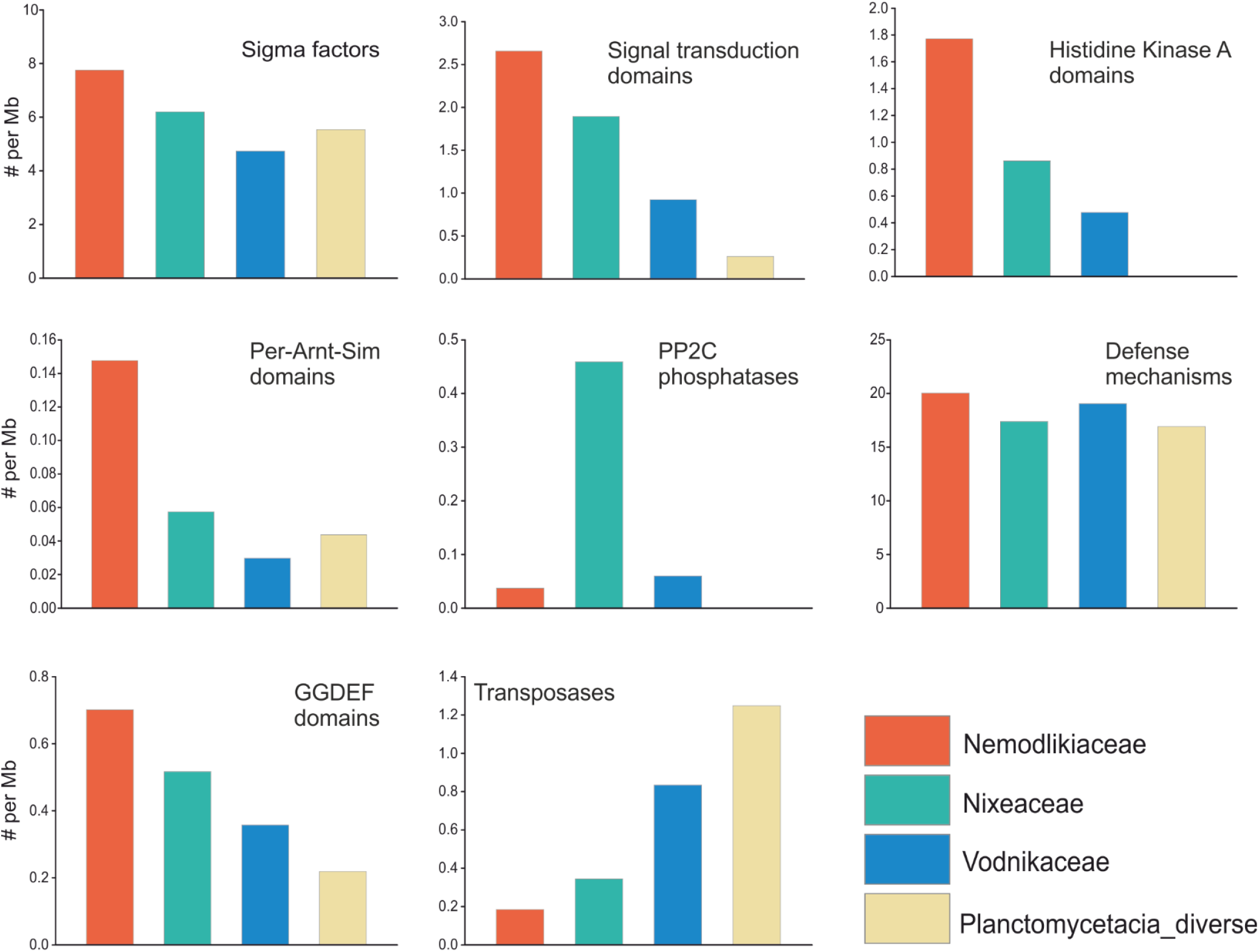
Abundance of signal transduction systems, defense mechanism and transposases in Nemodlikiaceae, Vodnikaceae, Nixeaceae and Planctomycetacia_diverse.

**Supplementary Figure 7.**
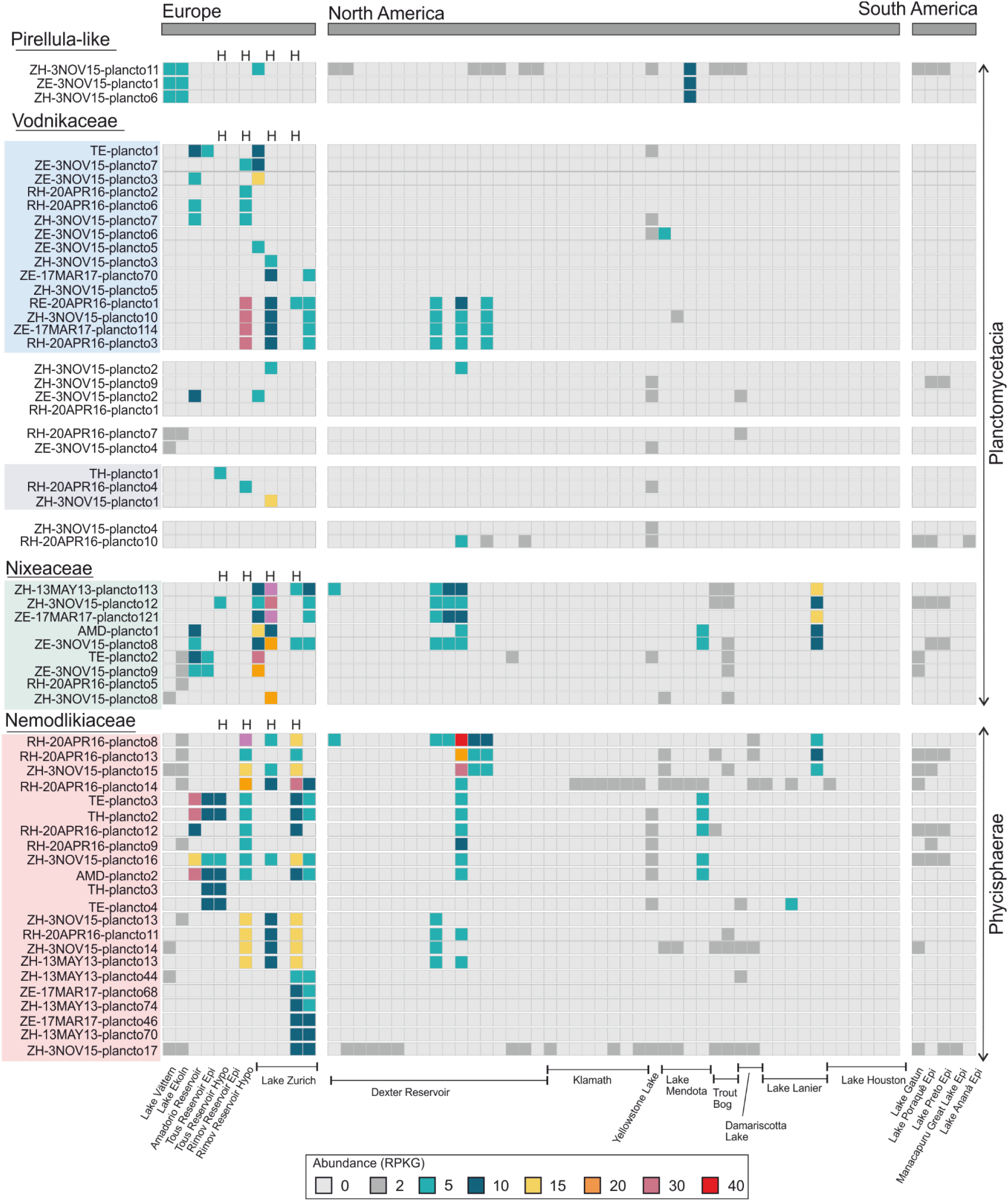
Heat-map of MAGs abundance (expressed as RPKG values) in 64 lacustrine datasets. The left side of the heat-map shows the MAGs grouped by phylogeny, while the bottom part of the figure shows the geographic distribution of the sample collection sites. Four hypolimnion datasets (Tous, Rimov 1 each, 2 from Lake Zurich)areindicatedwithan H.

**Supplementary Figure 8.**
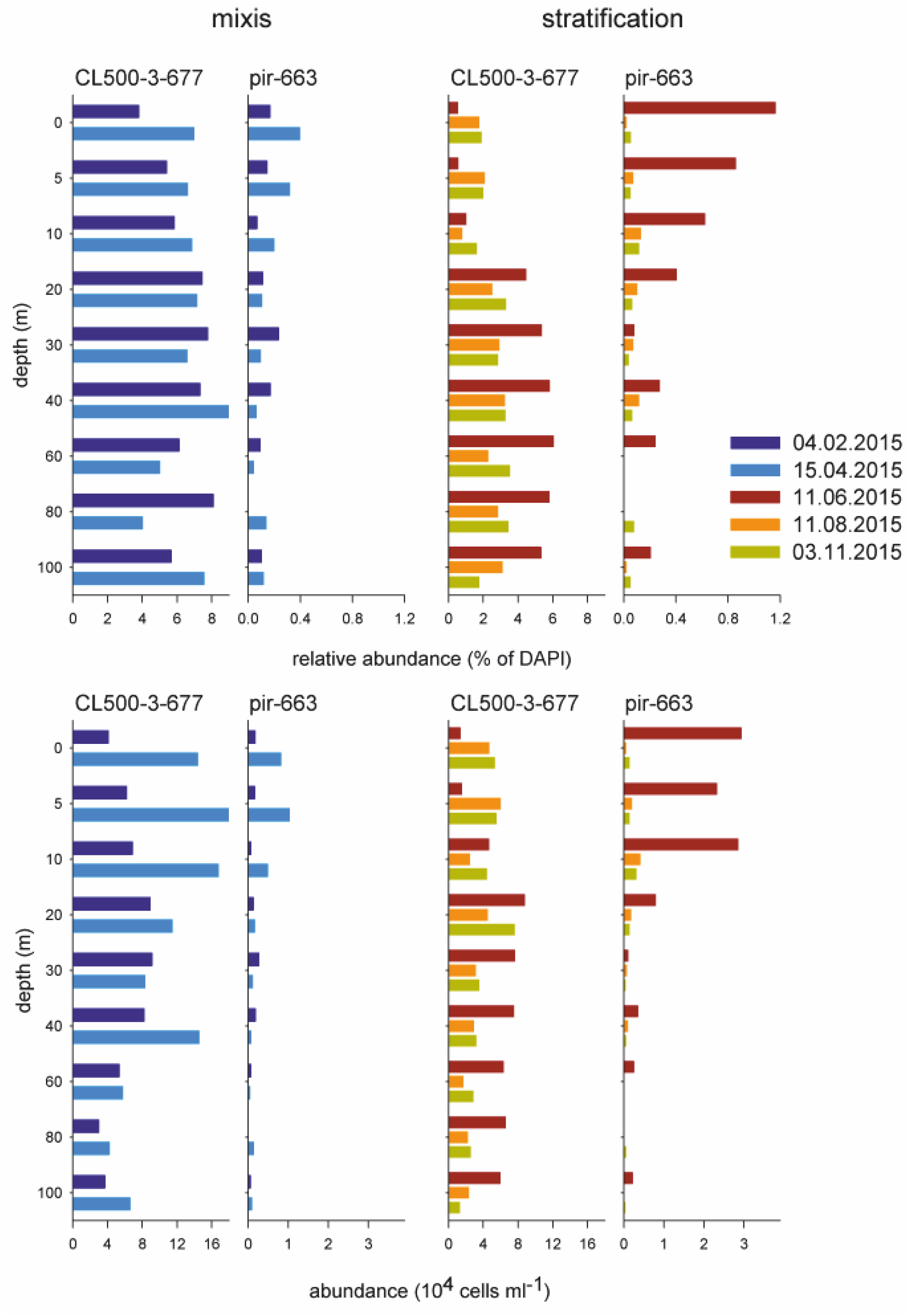
Vertical profiles of CARD-FISH abundances (top: relative, bottom: absolute) of two lineages of Planctomycetes in Lake Zurich during two different sampling periods in 2015 (mixis and stratification).

**Supplementary Figure 9.**
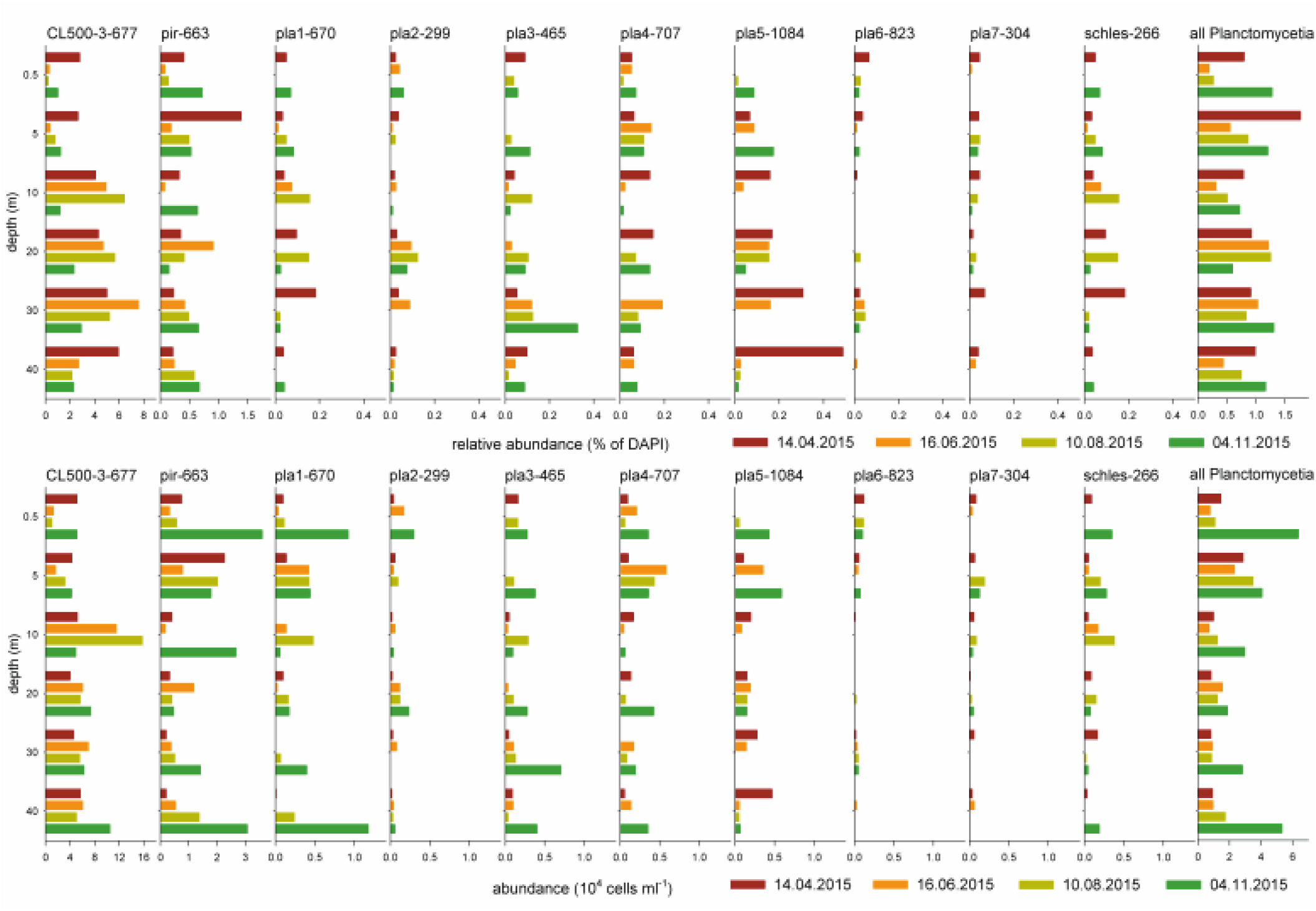
Vertical profiles of CARD-FISH abundances (top: relative, bottom: absolute) often lineages of Planctomycetes In the Rimov Reservoir during four different samplings in 2015.

**Supplementary Figure 10.**
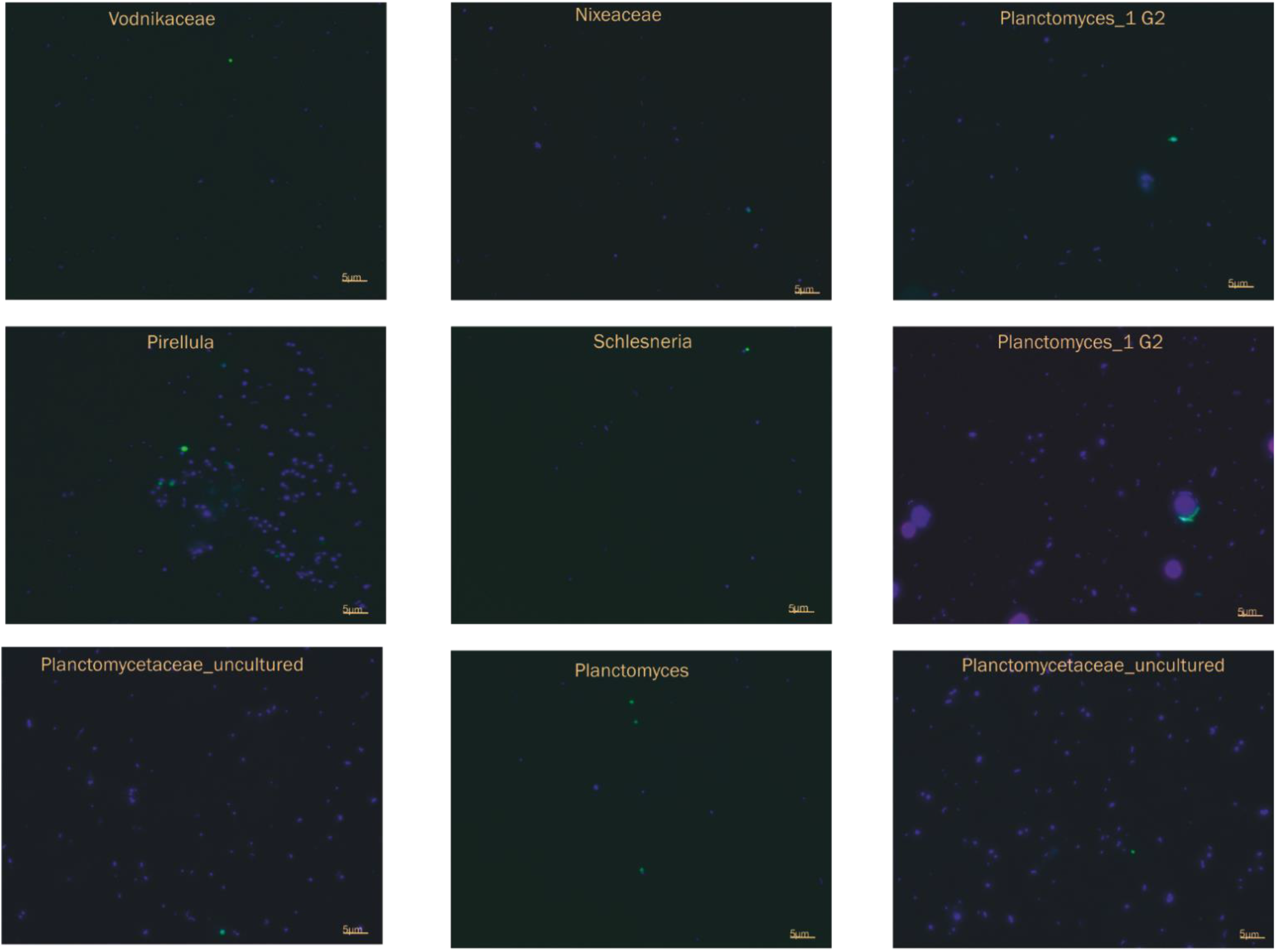
Superimposed images of CARD-FISH-stained Planctomycetes (green color) and DAPI-stained prokaryotes (blue color). The names of the panels are in accordance with Supplementary Figure 4. The scale bar is 5 μm.

**Supplementary Figure 11.**
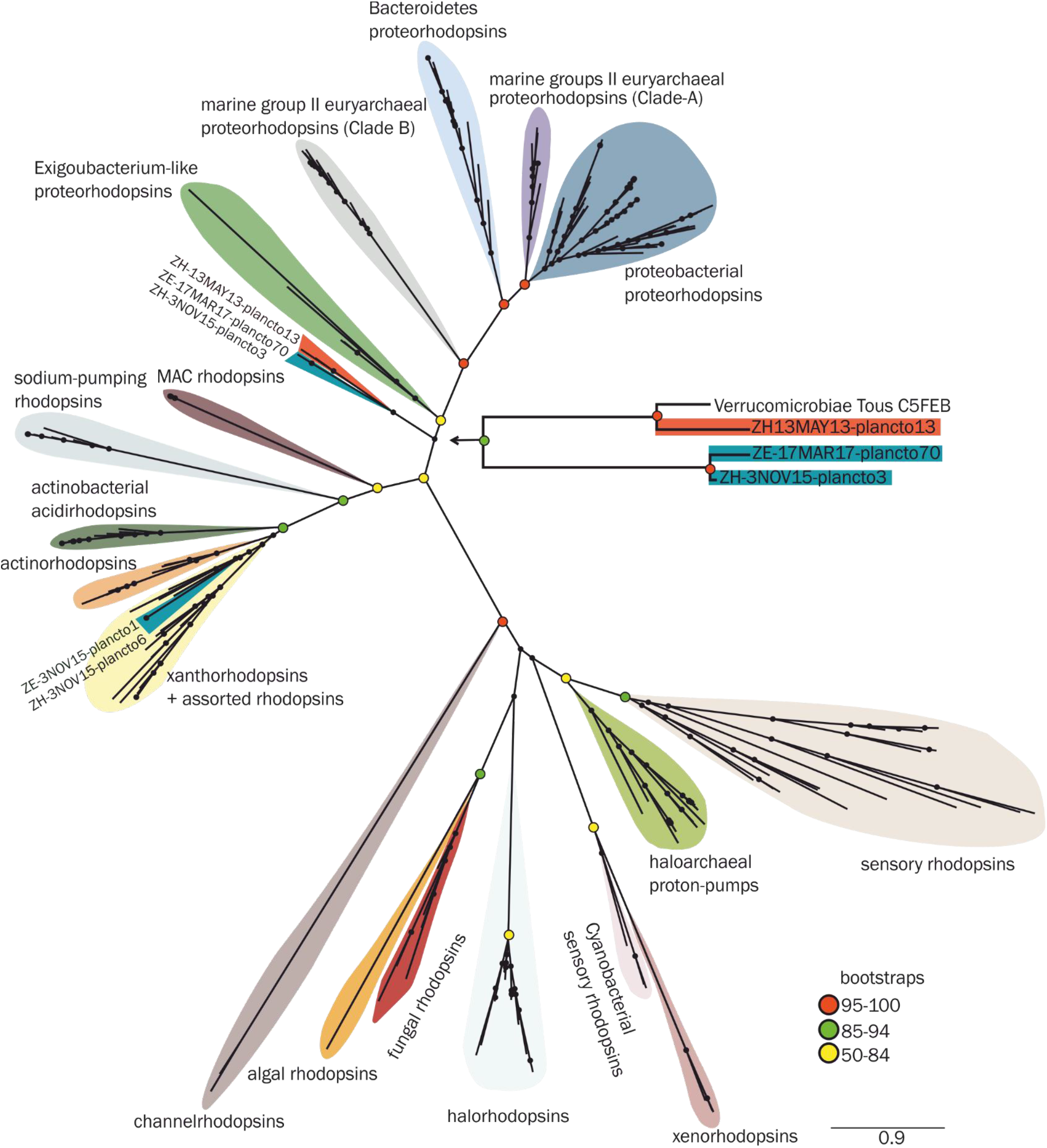
Maximum likelihood phylogenetic tree constructed using 254 rhodopsin sequences derived from diverse environments (freshwater, brackish, marine and hypersaline). The sequences belonging to freshwater MAGs are depicted in a subtree on the right side of the main figure.

**Supplementary Figure 12.**
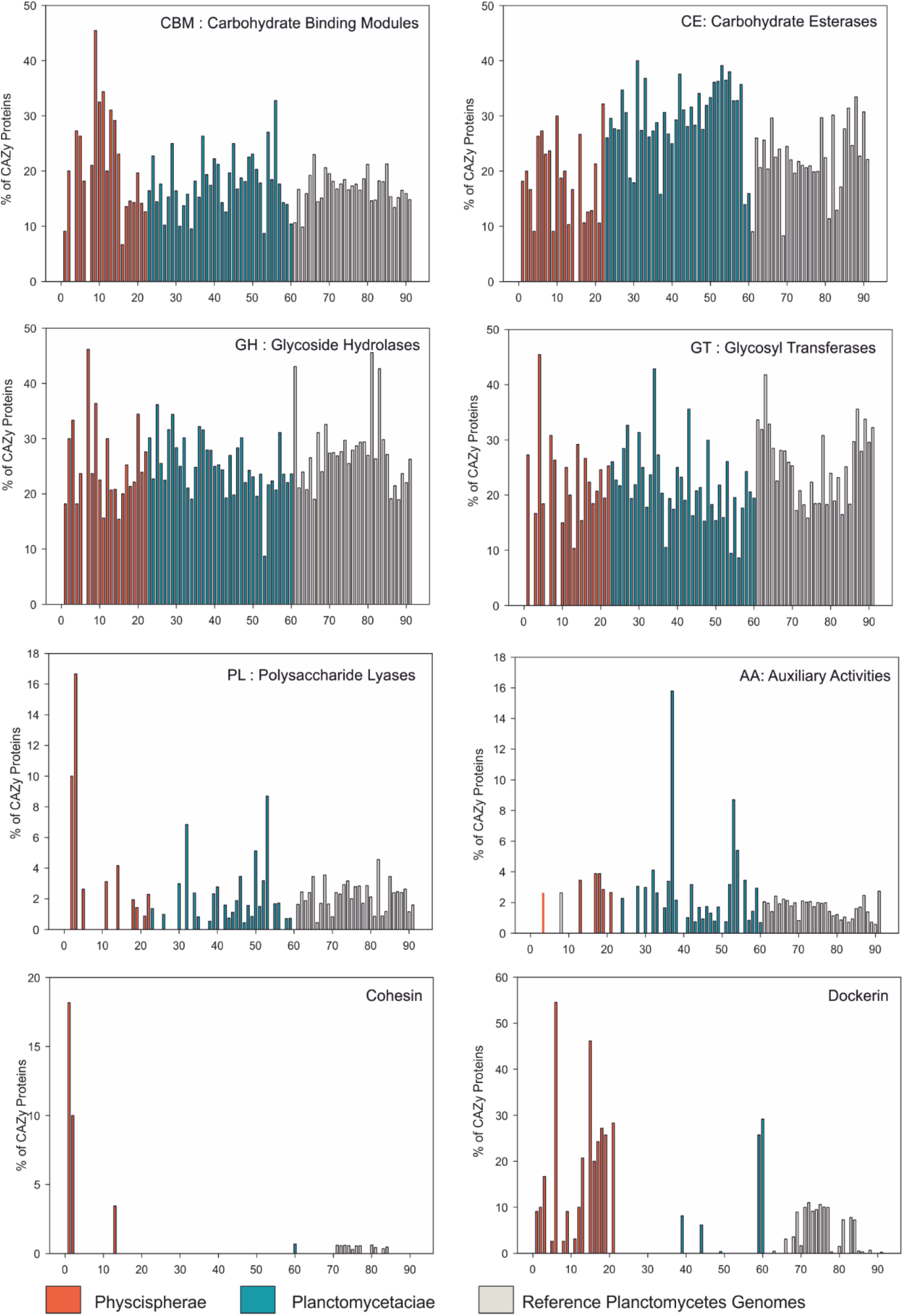
Families of structurally-related catalytic and carbohydrate-binding modules (CAZy proteins) found in freshwater Planctomycetes MAGs and reference genomes.

**Supplementary Figure 13.**
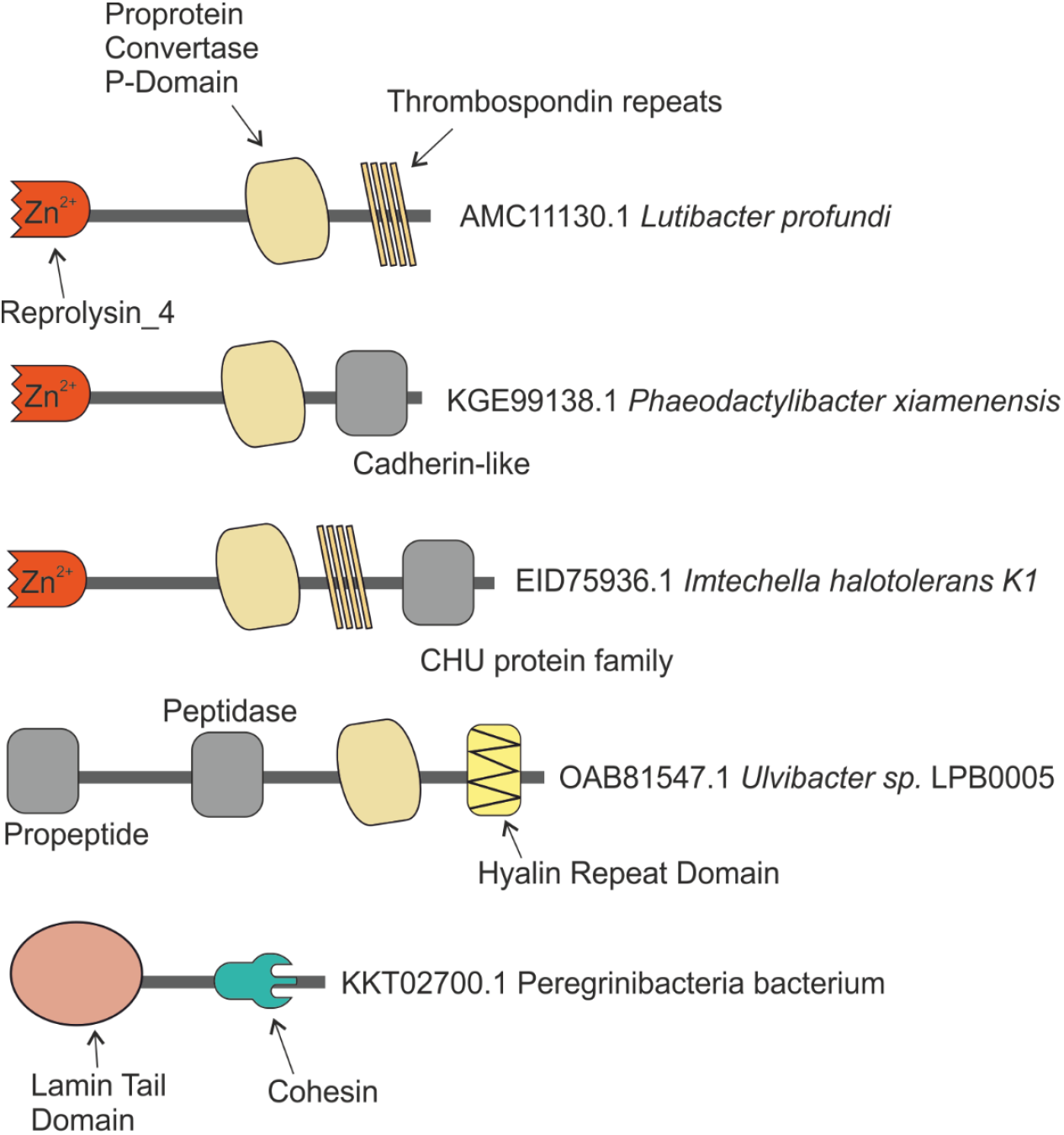
Planctosome protein domains (shown in colors) found in proteins belonging to diverse bacterial species. Other domains are shown in grey.

## Supplementary Materials and Methods

### Sampling and Sequencing

The meso-eutrophic Římov Reservoir (470 m a.s.l, 48°50’N, 14°29’E, Czech Republic) is a canyon-shaped dimictic water body with an area of 2.06 km^2^ (length 13.5 km, volume of 34.5 × 10^6^ m^3^, mean retention time 77 days, maximum depth 43 m), that was built during 1974–1979 by damming a 13.5 km long section of the River Malše [1]. The sampling was performed during Spring 2016 (April 20), above the deepest point of the reservoir by using a Friedinger sampler. Two multi-parametric probes were deployed in order to profile the physicochemical characteristics of the water column (temperature, pH, oxygen,; GRYF XBQ4, Havlíčkův Broc, CZ) and chlorophyll *a* (FluoroProbe TS-16-12, bbe Moldaenke, Kiel, Germany). 10 L of water were collected from 0.5 and 30 m depths and subjected to sequential peristaltic filtration through a series of 20, 5 and 0.2-μm-pore-size polycarbonate membrane filters (ø 142 mm) (Sterlitech Corporation, USA). The DNA was extracted from the 0.2 – 5-μm fraction, as described elsewhere [2] and subjected to deep shotgun sequencing (paired end, 150bp) on Illumina’s HiSeq 4 000 platform (BGI, Hong Kong).

The oligomesotrophic Lake Zurich (406 m a.s.l, 47°18’N, 8°34’E, Switzerland) is a perialpine, monomictic water body, with an area of 67.3 km^2^ (length 40 km, volume 3.3 km^3^, mean retention time 1.4 years, maximum depth 136 m). The sampling was conducted during an ongoing fortnightly monitoring program at the deepest point of the lake [3]. Vertical profiles of temperature, conductivity, turbidity, and oxygen were recorded with a YSI multiprobe (Yellow Springs Instruments, model 6 600) and the chlorophyll *a* concentration was measured with a submersible fluorescence probe (FluoroProbe TS-16-12, bbe Moldaenke, Kiel, Germany). Water samples from the following depths were collected with a Friedinger sampler and processed for sequencing: 5 and 80 m (13^th^ May 2013), 5 and 80 m (3^rd^ November 2015) and 2 m (17^th^ March 2017), respectively. Approx. 1-2 L of water was sequentially filtered onto 5 and 0.2 μm filters, and the genomic DNA was extracted from the 0.2 μm filter one by using the PowerBiofilm DNA Isolation Kit (Mo Bio Laboratories, Carlsbad, CA, USA). Library preparation of 550-bp fragments was done with a KAPA Hyper Prep Kit (Kapa Biosystems, Wilmington, MA, USA) and deep metagenomic sequencing (paired-end, 150 bp) was carried out on a HiSeq 2 000 instrument at the Functional Genomics Center Zurich.

### Classification of shotgun 16S rRNA gene fragments

FASTQ files (recovered from 298 environmental metagenomes: 64 lacustrine, 36 fluvial, 158 marine and 40 freshwater sediments, Extended Data) containing aquatic/sediment-derived raw shotgun reads, produced by second-generation sequencing platforms, were quality-filtered by a combination of bbduk.sh (adapter trimming and contaminant filtering) [4], bbmerge.sh *(de novo* adapter identification) [5] and sickle (quality trimming) [6]. Subsequently, they were converted to FASTA format and subsampled to 10 million sequences using reformat.sh [7]. These subsets (containing 10 million sequences each) were screened to identify RNA-like sequences by using UBLAST [8] against a non-redundant version of RDP database [9], that was previously clustered at 85% sequence identity by UCLUST [8] and contained 7 552 sequences with a length ≥800 bp. The sequences that matched the RDP database at an E-value <1e-5 were considered candidate 16S rRNA gene sequences and screened using SSU-ALIGN [10]. The *bona fide* 16S rRNA gene sequences (as identified by SSU-ALIGN) were further compared by BLAST [11], in nucleotide space (using as cut-off the E-value 1e-5), against a curated SILVA SSU database [12] that contained 447 012 sequences, and classified if the sequence identity was ≥80% and the alignment length was ?90 bp (sequences failing these thresholds were not used for downstream analyses).

### Assembly and binning

Ten lacustrine shotgun metagenomic datasets generated from lakes with contrasting trophic states (i.e. Lake Zurich and Římov, Tous and Amadorio reservoirs) were used for in-depth analyses. The metagenomic datasets derived from the Spanish freshwater reservoirs (i.e. Tous and Amadorio) were recovered from NCBI’s SRA database, under the accession numbers SRR1173821 (Amadorio), SRR4198666 and SRR4198832 (Tous). Seven shotgun metagenomic libraries, generated from Římov Reservoir (n=2) and Lake Zurich (n=5), were sequenced during this study. All raw metagenomic sequences were filtered to remove low quality bases/reads as mentioned above, by using a combination of bbduk.sh [4], bbmerge.sh [5] and sickle [6] (Římov Reservoir datasets) or trimmomatic [13] (Lake Zurich datasets). The obtained high quality sequences were then assembled independently with MEGAHIT v1.1.1 [14] using the parameters: --min-count 2 and --k-step 10 (k-mer range was 31 – 99 for the Tous and Amadorio datasets, and 31 – 149 for the Římov datasets) or metaSPAdes [15] (k-mer range 21 – 127) for the Lake Zurich datasets.

For generating Planctomycetes metagenome-assembled genomes (MAGs), we used a combination of taxonomy-dependent and -independent binning techniques. Firstly, a supervised alignment-based method (e.g. taxonomic assignment according to the best hits) was used on the assembled contigs that had a minimum length of 5 kbp. Briefly, their protein coding sequences predicted by MetaProdigal [16] were annotated using UBLAST [8] (with the lenient cut-offs: E-value 1e-3, similarity 10%, coverage 10%, bitscore 50) against an in-house curated Prokaryotic RefSeq Genomes Release 82 database, that was amended by addition of supplementary genomes. The revision of RefSeq database was done by including all Planctomycetes genomes (including MAGs and SAGs) publicly available in NCBI Genome database (i.e. 102 entries in May 2017). The genomes that were affiliated to Planctomycetes phylum (as predicted by PhyloPhlAn) and had a genome completeness (as estimated by CheckM) [17] higher than 10% were merged with the above-mentioned RefSeq release. Prior to inclusion in RefSeq, 13 genomes (that lacked annotation) were annotated using Prokka [18]. Secondly, we selected from all the assembled metagenomic datasets (i.e. 10 datasets) the contigs with a minimum length of 5 kbp that gave more than 50% best UBLAST hits to Planctomycetes (if from the total number of proteins present in one contig more than 50% gave hits to Planctomycetes we considered the contig belonging to Planctomycetes phylum) and used them further for taxonomy–independent binning. For this step, we used the contigs’ mean base coverage, computed by bbwrap.sh (with default parameters) [19], to perform hybrid binning using tetranucleotide frequencies and abundance data *via* MetaBAT [20] (using the presets –superspecific and --minCorr 99). Prior to downstream analyses, the bins that were found to be poorly resolved (i.e. have more than 10% redundancy) were further refined using anvi’o software [21] as described elsewhere (http://merenlab.org/2016/06/22/anvio-tutorial-v2/). The obtained bins that were found to be taxonomically affiliated with Planctomycetes (as established by PhyloPhlAn) and to have a genome completeness higher than 10% (as determined by using 360 Planctomycetes marker genes in CheckM) were denominated as Planctomycetes MAGs.

### Phylogenomics

In order to investigate if the obtained 60 Planctomycetes MAGs were identical to previously described ones, we performed genome distance estimations, using Mash software [22] (with the parameters k-mer 25 and sketch size 5 000), against the Planctomycetes genomes publicly available in NCBI Genome database (102 entries in May 2017).

The average MAG coverage depth (defined as the average number of reads covering a base pair in the reference MAG) was computed by using BBMap version 36.19 (with default settings) [23] and quality-trimmed metagenomic reads. In order to estimate the abundance of each MAG within and between metagenomes, we calculated RPKG values (i.e. the number of reads recruited per kilobase of genome per gigabase of metagenome) using an in-house pipeline. Briefly, in order to avoid analysis bias we concatenated the contigs belonging to each MAG and masked all the rRNA gene sequences present. Subsequently, BLASTN [11] (with the cutoffs: alignment length ≥50 nt, identity > 95%, E-value <= 1e-5) was used in order to align the quality filtered shotgun reads (20 million reads each from 64 freshwater metagenomic datasets) against the 60 Planctomycetes MAGs. The obtained BLAST best-hits results were further used to compute RPKG values. In order to assess the genetic diversity of the Planctomycetes populations, we used blast-tools [24] to plot the best-hits results generated by performing BLASTN (with 180 million sequences equally sub-sampled from 10 metagenomes generated from Lake Zurich and, Římov, Tous and Amadorio reservoirs) against the 60 MAGs.

In order to establish the evolutionary relationships among the 60 MAGs (with variable degrees of genome completeness) and previously available Planctomycetes genomes (in NCBI Genome repository), we carried out a phylogenomic analysis using PhyloPhlAn [25]. Briefly, the CDSs predicted in Prodigal’s metagenomic mode [16] were translated to protein sequences and screened for the presence of 400 universally conserved and phylogenetically discriminating proteins (found in PhyloPhlAn database) by USEARCH [8] (E-value < 1e-40). The minimum number of proteins used was 35 (for ZH-3NOV15-plancto17), the maximum 319 (RH-20APR16-plancto1) and the median 170. The homologs of these proteins were independently aligned by MUSCLE [26], concatenated and further used in generating a maximum likelihood tree with FastTree software (JTT+CAT model) [27]. Subtrees were constructed by concatenating and aligning conserved proteins as describes elsewhere [28].

The average amino acid identity (AAI) within coherent phylogenomic groups was determined by performing whole-genome pairwise CDSs comparisons, using BLAST software, as previously described by Konstantinidis and Tiedje [29]. Taxonomic categories for the MAGs where defined using the standards suggested by Konstantinidis *et al.* [30].

Planctomycetes *in situ* replication rates were determined based on measuring the rate of the decrease in average sequence coverage across all genomic fragments (present in one MAG), by using iRep software [24]. Briefly, quality-filtered shotgun reads were mapped against the MAGs (>=75% complete, <=175 fragments/Mbp sequence, and <=4% contamination) recovered from the same metagenome by Bowtie 2 (version 2.3.4) [31] with --very-sensitive option. The obtained mapping files, in SAM format, were used for calculating an index of replication (iRep) based on the sequencing coverage trend that results from bidirectional genome replication from a single point of origin as described by Brown *et al.* [24]. Genome Annotation

MAGs *de novo* gene predictions were performed by Prokka [18]. BlastKOALA [32] was used to assign KO identifiers (K numbers) to orthologous genes present in the 60 MAGs. The K numbers were further mapped to KEGG pathways, BRITE hierarchies, and KEGG modules for inferring the systemic functions of individual MAGS. The annotations were further refined by using the standard operation procedures from the Rapid Annotations using Subsystems Technology server [33]. Additional gene annotations were performed by protein sequence searches (using hmmscan with E-value 1e-5) [34] against the HMM databases COG [35] and TIGRfam [36]. The carbohydrate-active enzymes were annotated using the dbCAN-seq database [37]. Several protein sequences were further analyzed using jackhmmer [38] and Phyre2 [39].

### Phylogenetics

The 16S rRNA gene sequences present in the MAGs were identified by SSU-ALIGN, aligned by SINA (https://www.arb-silva.de/aligner/), imported in ARB software [40] using the SILVA SSU Ref 123 database and manual refinements of alignments, and used for the construction of a RAxML [41] maximum likelihood tree (100 bootstraps, GTRGAMMA model). The rhodopsin sequences identified by HMMER [34] were aligned with MAFFT under L-INS-i model [42], and used for a maximum likelihood tree construction (100 bootstraps) with FastTree2 [27].

### Probe design and CARD-FISH

The 16S rRNA gene sequences present in MAGs as well as 16S rRNA sequences extracted from the raw metagenomics reads were used for probe design for fluorescence *in situ* hybridization followed by catalyzed reporter deposition (CARD-FISH). The RAxML tree (Supplementary Figure S4) served as backbone, and probes for 10 monophyletic lineages containing MAG sequences or high amounts of sequences extracted from reads were subjected to probe design using the probe_desing and probe_check tools in ARB [40]. The resulting probes were checked *in silico* using mathFISH [43] and in the laboratory with different formamide concentrations until stringent hybridization conditions were achieved. CARD-FISH was done for 28 samples from Římov Reservoir and 45 samples from Lake Zurich collected in 2015: Římov Reservoir was sampled during the spring phytoplankton bloom (April 14^th^), early summer (June 16^th^), late summer (August 10^th^), and autumn (November 04^th^) at 0, 5, 10, 20, 30, and 40m depths and Lake Zurich was sampled during winter mixis (February 4^th^), the spring phytoplankton bloom (April 15^th^), early summer (June 11^th^), late summer (August 11^th^), and autumn (November 3^th^) at 0, 5, 10, 20, 30, 40, 60, 80, and 100m depths. CARD-FISH was done with fluorescein-labeled tyramides as previously described [44] and analyzed by fully automated high-throughput microscopy [45]. Interfering autofluorescent cyanobacteria or debris particle were individually excluded from hybridized cells and at least 10 high quality images or >1000 DAPI stained bacteria were analyzed per sample. Micrographs of CARD-FISH stained Planctomycetes lineages were recorded with a highly sensitive charge-coupled device (CCD) camera (Vosskühler) at a magnification of 1000 x and cell sizes were estimated using the software LUCIA (Laboratory Imaging Prague, Czech Republic) following a previously described workflow [46].

### Accession numbers

All sequence data produced during the study is deposited in the Sequence Read Archive (SRA) database of the National Center for Biotechnology Information (NCBI) and could be found linked to the Bioprojects PRJNA429141 (Řimov Reservoir) and PRJNA428721 (Lake Zurich). All MAGs used in this study can be accessed under the Bioproject PRJNA449258.

